# High-energy demand and nutrient exhaustion in MTCH2 knockout cells

**DOI:** 10.1101/2023.12.15.571941

**Authors:** Sabita Chourasia, Christopher Petucci, Hu Wang, Xianlin Han, Ehud Sivan, Alexander Brandis, Tevie Mehlman, Sergey Malitsky, Maxim Itkin, Ron Rotkopf, Limor Regev, Yehudit Zaltsman, Atan Gross

## Abstract

Mitochondrial carrier homolog 2 (MTCH2) is a regulator of apoptosis, mitochondrial dynamics, and metabolism. Loss of MTCH2 results in mitochondrial fragmentation, an increase in whole-body energy utilization, and protection from diet-induced obesity. We now show using temporal metabolomics that MTCH2 deletion results in a high ATP demand, an oxidized environment, a high lipid/amino acid/carbohydrate metabolism, and in the decrease of many metabolites. Lipidomics analyses show a strategic adaptive decrease in membrane lipids and an increase in storage lipids in MTCH2 knockout cells. Importantly, all the metabolic changes in the MTCH2 knockout cells were rescued by MTCH2 re-expression. Interestingly, this imbalance in energy metabolism and reductive potential triggered by MTCH2-deletion inhibits adipocyte differentiation, an energy consuming reductive biosynthetic process. In summary, loss of MTCH2 results in an increase in energy demand that triggers a catabolic and oxidizing environment, which fails to fuel the anabolic processes during adipocyte differentiation.

## Introduction

Energy homeostasis is a fundamental physiological process crucial for the survival and well-being of organisms. The energy homeostasis of the organism is fine-tuned by dynamic processes that maintain the balance between energy intake and expenditure^1,2^. AMP-activated protein kinase (AMPK), sirtuins (such as SIRT1) and mTOR, act as cellular energy sensors, influencing metabolic pathways and promoting energy conservation during low energy states^3–6^.

At the core of this intricate system lies the mitochondria that plays a pivotal role in energy production, sensing, adapting, and responding to the cellular energy demands. Mitochondria are the primary sites of cellular respiration, where the process of oxidative phosphorylation (OXPHOS) converts nutrients into adenosine triphosphate (ATP)^7–10^. Redox cofactors, the oxidized (NAD^+^) and reduced (NADH) forms of nicotinamide, act as “fuel” for the mitochondria^11–13^. The mitochondrial NAD^+^/NADH pool are substrates for OXPHOS, where 90% of cellular ATP production takes place. Along with ADP^13^, NAD^+^ also plays an important role in the regulation of the Krebs cycle. The mitochondrial-redox cofactor/fuel relationship dictates the metabolic status and sets the entire blueprint for cellular bioenergetics and cell behavior^14^.

Mitochondria are dynamic entities: they undergo constant fission and fusion processes known as mitochondrial dynamics. This dynamic behavior is essential for maintaining a healthy mitochondrial population, responding to energy demands, and orchestrating adaptability in the face of changing metabolic states^8,9,15,16^. In addition, mitochondria biogenesis is intricately linked to energy homeostasis. Peroxisome proliferator-activated receptor gamma coactivator 1-alpha (PGC-1α) is a master regulator that coordinates mitochondrial biogenesis in response to energy demand, environmental cues, and cellular stress^17,18^. Mitochondria maintain their health by a delicate balance between biogenesis of new mitochondria and clearance of old and dysfunctional mitochondria by mitophagy^19^.

Metabolism plays a crucial role in cell differentiation and adaptation during cellular energy stress. The dynamic shift from glycolysis to OXPHOS during cellular differentiation underscores the adaptability and plasticity of cellular metabolism. Glucose metabolism is pivotal for sustaining the energy demands of undifferentiated and proliferating cells (Warburg effect)^20^. The glycolysis- to-OXPHOS switch stands at the crossroads of metabolism and cell fate, revealing the intimate connection between energy metabolism and cellular differentiation^21^. Adipocyte differentiation, or adipogenesis, involves a finely-tuned balance of energy utilization, demanding dynamic shifts in energy metabolism to support lipid accumulation and storage^22^. Transcriptional regulators, including PPARγ, C/EBPs, and AMPK, act as key nodes governing the balance between energy storage and expenditure during adipocyte differentiation. Their intricate interplay shapes the metabolic landscape of adipose tissue^23^. NAD^+^ acts as a critical signaling molecule for activation of PPARγ and C/EBPs during adipocyte differentiation^24^. During adipocyte differentiation, the cytoplasmic NAD^+^ pool is engaged in meeting cellular metabolic demand by regulating glucose metabolism and the nuclear pool involved in gene regulation^25,26^.

Mitochondrial carrier homologue 2 (MTCH2; also named MIMP or SLC25A50), stands out as a unique member of the mitochondrial carrier protein family, positioned at the outer mitochondrial membrane (OMM)^27^. Initially acknowledged for its role in mediating apoptosis^28,29^ subsequent studies uncovered its multifaceted involvement in regulating mitochondria/whole-body metabolism and hematopoietic stem cell fate^30,31^. In addition, multiple genome-wide association studies have associated the MTCH2 locus with metabolic disorders such as diabetes and obesity^32–36^. More recently, MTCH2 has been implicated in the regulation of mitochondrial dynamics, broadening its spectrum of cellular functions^37,38^. Various investigations, including genetic models in diverse organisms such as *C. elegans*, *Zebrafish*, and mice, have shed light on the involvement of MTCH2 in lipid metabolism^39–42^. MTCH2 deletion has been linked to diminished lipid synthesis and storage, underscoring its critical involvement in lipid homeostasis^30,40,42^. Conversely, increased MTCH2 expression has been associated with elevated lipid storage, highlighting its dynamic regulatory role in lipid metabolism^39^.

A recent study has illuminated the contribution of MTCH2 to mitochondrial fusion, establishing a connection between MTCH2 and the pro-mitochondrial fusion lipid lysophosphatidic acid (LPA)^38^. This study proposed that MTCH2 plays a pivotal role in the biogenesis and transfer of lipids from the ER to the mitochondria, and that this transfer plays a critical role in mitochondria fusion. In addition, MTCH2 was recently demonstrated to act as a protein insertase^43^, and possibly also as a scramblase^44,45^.

In our present paper, we show that loss of MTCH2 results in increased energy demand and increased metabolism, across many pathways, most likely to satisfy this demand.

## Results

### A high ATP demand and an oxidizing environment in MTCH2 knockout cells

To assess MTCH2’s metabolic function, we performed temporal targeted metabolomics analyses. The analyses were performed on HeLa stable cell lines, six clones from each genotype: wild type (WT) with empty vector, MTCH2 knockout (MKO), and MKO reconstituted with MTCH2 (MKO-R). All stable lines were cultured in complete media (CM), left to grow overnight, then the media was changed (considered as time 0) and samples were taken at 1, 6, 12, and 18-hrs post media change. Hierarchical clustering and principal component analysis (PCA) based on the differential levels of metabolites between the three groups showed that while WT and MKO-R cells are clustered together, the MKO cells are a distinct sub-population of cells (Fig. 1A). These results suggest that MTCH2 knockout results in a prominent metabolic change, which is largely rescued to the WT metabolic state by re-introducing MTCH2.

**Figure 1.**
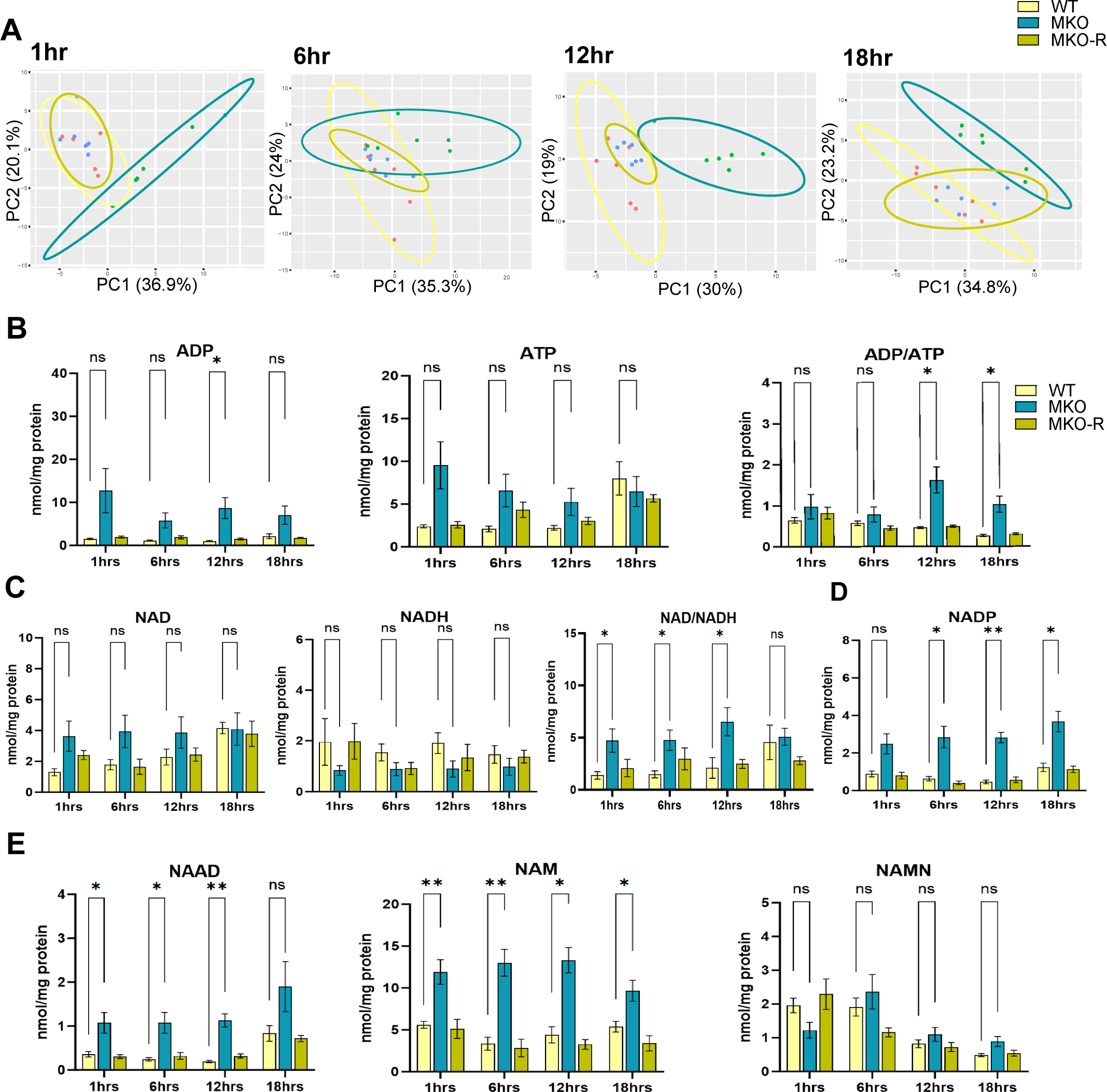
A high ATP demand and an oxidizing environment in MTCH2 knockout cells. A. PCA plots of WT, MKO and MKO-R cell lines. Ellipses describe 95% confidence intervals. B. Average levels of ADP, ATP, and ADP/ATP ratios in all 3 cell lines at all four time points. C. Average levels of NAD^+^, NADH, and NAD^+^/NADH ratios in all 3 cell lines at all four time points. D. Average levels of NADP^+^ in all 3 cell lines in all four time points. E. Average levels of NAAD, NAM, and NAMN in all 3 cell lines at all four time points. Results in all graphs in A-E are presented as mean ± SEM (\**p*<0.05, \*\**p*<0.001; two-way ANOVA with Dunnett multiple comparison test; n=6 biological replicates).

Detailed analysis of the results revealed that the MKO cells showed: 1) Trend increases in ADP and ATP levels resulting in an increase in the ADP/ATP ratio (Fig. 1B). 2) Trend increases in NAD^+^ and a trend decrease in NADH levels resulting in an increase in the NAD^+^/NADH ratio (Fig. 1C). 3) An increase in NADP^+^ levels (Fig. 1D). 4) An increase in the nicotinamide precursors nicotinamide adenine dinucleotide (NAAD) and nicotinamide (NAM), and a trend decrease in nicotinamide mononucleotide (NAMN) (Fig. 1E), which point to an overall change in nicotinamide metabolism.

Importantly, the majority of the metabolic changes measured in the MKO cells were rescued to WT levels in the MKO-R cells (Fig. 1B-E), suggesting that the observed metabolic changes were due to MTCH2 knockout. An increase in the ADP/ATP ratio stimulates OXPHOS; an increase in the NAD^+^/NADH ratio and NADP^+^ levels represent an oxidized environment, which leads to stimulation of glycolysis^46,47^. Collectively, these results suggest that MTCH2 knockout results in the stimulation of oxidative metabolism and ATP production to meet the increased cellular demand for ATP.

### An increase in amino acid/lipid/carbohydrate metabolism and a decrease in many metabolites in MKO cells

The metabolomics analyses revealed additional important changes in many more nutrient substrates, which included a decrease in most amino acids (Fig. 2A and Fig. S2A). Notably, the most significant change was seen in glutamine (Fig. 2A, left top graph), one of the major amino acid-nutrient sources^48^. A decrease in amino acids usually represents an increase in TCA cycle metabolism^49^. Indeed, we found a decrease of TCA cycle intermediates 18hr post media change in the MKO cells (Fig. S2B).

**Figure 2.**
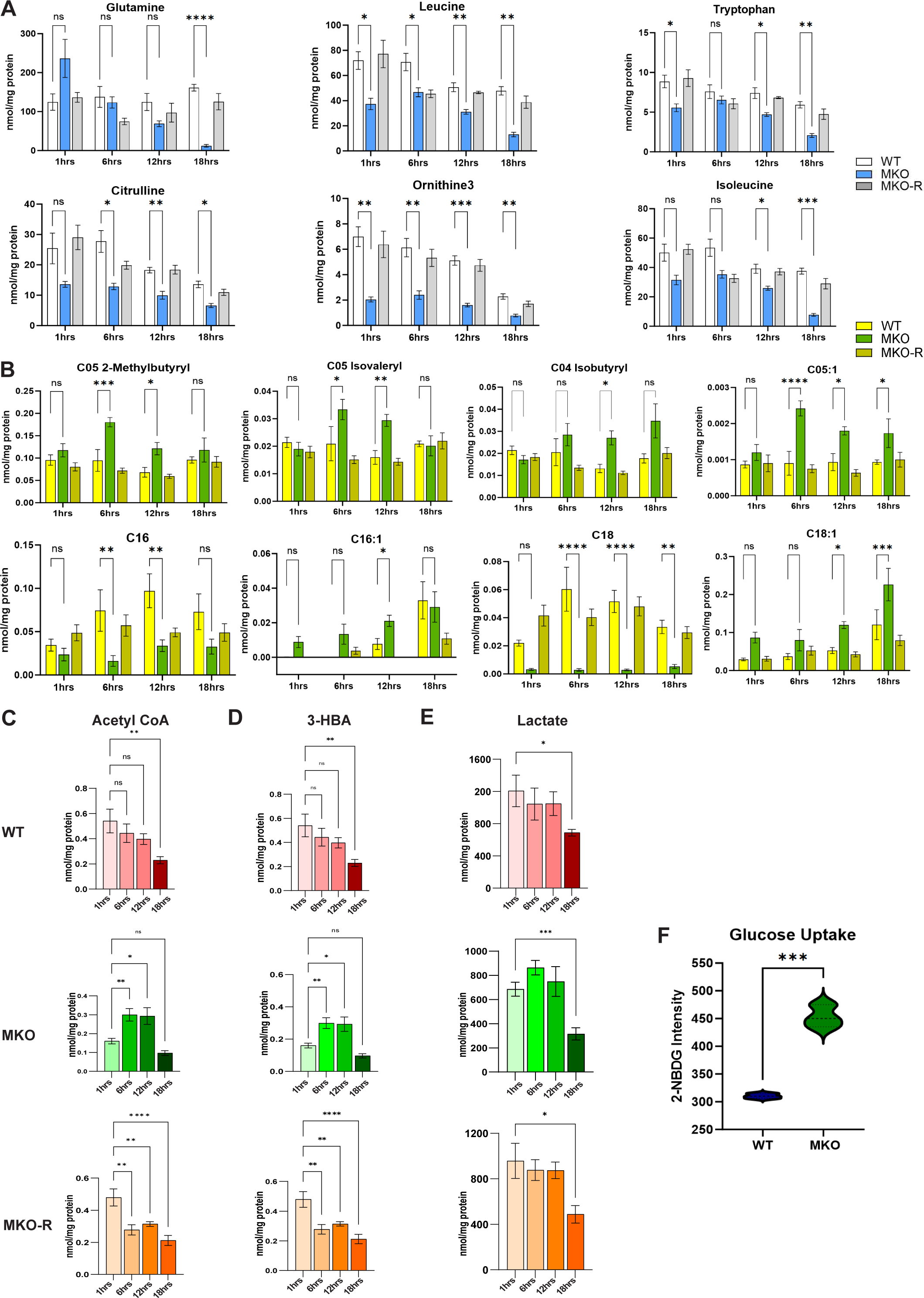
An increase in amino acid/lipid/carbohydrate metabolism in MKO cells. A. Average levels of a set of amino acids in all 3 cell lines at all four time points. B. Average levels of acyl carnitines in all 3 cell lines at all four time points. C, D, E. Average levels of acetyl CoA (C), 3-HBA (D) and lactate (E) in all 3 cell lines at all four time points. F. Glucose uptake from media of WT and MKO cells after a two-hour glucose starvation. Results in all graphs in A-F are presented as mean ± SEM (\**p*<0.05, \*\**p*<0.001, \*\*\**p*<0.0003, \*\*\*\**p*<0.0007; two-way ANOVA with Dunnett multiple comparison test) (In A-E: n=6 biological replicates; In F: n=3 technical replicates).

Acyl carnitines are a “readymade” form of fatty acids, which can enter mitochondria for breakdown. Overall, 22 species of acyl carnitines were detected in our metabolomics analyses and among them three were branched chain amino acid (BCAAs)-derived acyl carnitines (C05-2-Methylbutyryl, C05-Isovaleryl, and C04-Isobutyryl), five unsaturated acyl carnitines (C05:1, C16:1, C18:1, C14:2, C22:5) and eight saturated acyl carnitines (Fig. 2B and Fig. S2C). The MKO cells showed a sequential increase in BCAAs and unsaturated short chain (C05:1) acyl carnitines at 6- and 12-hrs post media change and an increase in part of the unsaturated acyl carnitines at later stages (Fig. 2B). Most changes in the MKO cells were rescued to WT levels in the MKO-R cells, again suggesting that the observed metabolic changes were due to MTCH2 knockout. The increase in acyl carnitines in the MKO cells is consistent with the idea of an increase in the transport and breakdown of fatty acids in mitochondria to meet the increase in cellular demand for ATP.

On the other hand, MKO cells show a decrease in part of the short and long chain saturated acyl carnitines that were rescued to WT levels in the MKO-R cells (Fig. S2C). An interesting comparison can be seen between two of the saturated acyl carnitines, C16 and C18, which their levels are decreased, whereas the levels of their two unsaturated counterpart forms, C16:1 and C18:1, were increased (Fig. 2B). Thus, the acyl carnitine profile suggests that 1- to 12-hrs post media change the MKO cells use BCAAs as a nutrient source, and later shift to unsaturated acyl carnitines, specifically to the C16:1 and C18:1 forms.

The levels of acetyl CoA, 3-hydroxybutyrate (3-HBA) and lactate also showed interesting differences. The main function of acetyl CoA is to deliver the acetyl group to the TCA cycle to be oxidized for energy production^50,51^. 1hr post media change, the levels of acetyl CoA were close to three-fold lower in the MKO cells as compared to WT and MKO-R cells (Fig. 2C). However, by 6- and 12hrs its levels in the MKO cells increased almost two-fold, whereas the levels in the WT and MKO-R cells gradually decreased (Fig. 2C). At 18hrs post media change the picture flipped back again, and the MKO cells showed lower levels of acetyl CoA as compared to the levels in the WT and MKO-R cells (Fig. 2C). These results suggest that there is higher metabolism of acetyl CoA in the MKO cells leading to a bell-shape dynamics (low-high-low levels).

A similar bell-shape dynamics was seen in the levels of the ketone body 3-hydroxybutyrate (3-HBA; Fig. 2D). 3-HBA is an alternative product of fatty acid oxidation and can be used as an alternate energy source in the absence of sufficient glucose or in preference towards lipogenic diet ^52,53^. A similar bell-shape dynamics was also seen in the levels of lactate (Fig. 2E), although the differences in lactate levels were less pronounced and did not reach statistical significance. In general, a more prominent increase in lactate means a more prominent severity of a condition^54,55^. The lower levels observed in acetyl CoA, 3-HBA and lactate 18hrs post media change in the MKO cells (Fig. 2C-E), led us to perform untargeted global metabolic profiling of cells 30hrs post media change. This profiling showed a prominent decrease of many metabolites in MKO cells (Fig. S2D and Table 1), metabolites involved in glycolysis, TCA cycle, pentose-phosphate pathway (PPP), and nucleotides. Most of these metabolic changes were largely rescued in the MKO-R cells. As might be expected from the results, the MKO cells showed a ∼1.5-fold higher uptake of glucose as compared to WT cells (Fig. 2F).

**Table 1.** Results of ANOVA per metabolite, followed by Dunnett’s post-hoc test to compare KO and Rescue to WT. Each contrast is described by log-fold change, p-value and fdr-corrected p-value (Fig 2SD-Continued).

Taken together, the results presented above are consistent with the idea that the increased amino acid/lipid/carbohydrate metabolism and substantial decrease of many metabolites in MKO cells is most likely due to their increased utilization to meet the increased cellular energy demand.

### Membrane lipids decrease and storage lipids and lipid droplets increase in MKO cells

Nutrient depletion leads to an adaptive process in which cells catabolize membrane lipids, manifested in fatty acids becoming available for energy storage (i.e., triglycerides (TAG)) or for energy production^56^. Targeted lipidomics on cells harvested 20hrs post media change, revealed a prominent decrease in the levels of many membrane lipids in MKO cells (Fig. 3A, left panel, and Table 2). Importantly, these changes in lipids were largely restored in the MKO-R cells (Fig. 3A, left and right panels). The MKO cells also showed a decrease in free fatty acids (FFA), non-esterified fatty acids (NEFA) (Fig. 3A, top right panel, and Fig. S3A). On the other hand, the level of esterified-fatty acids (TAG-fatty acid content) was increased in MKO cells (Fig. 3A, top right panel and Fig. S3B). A Volcano plot showed increased levels of the storage lipids TAG and cholesterol ester (CE) in MKO cells (Fig. 3A, bottom right panel). Interestingly, TAG is mostly composed of C16:0, C16:1 and C18:1 fatty acids, and two of these three were increased in the MKO cells (Fig. 2B).

**Figure 3.**
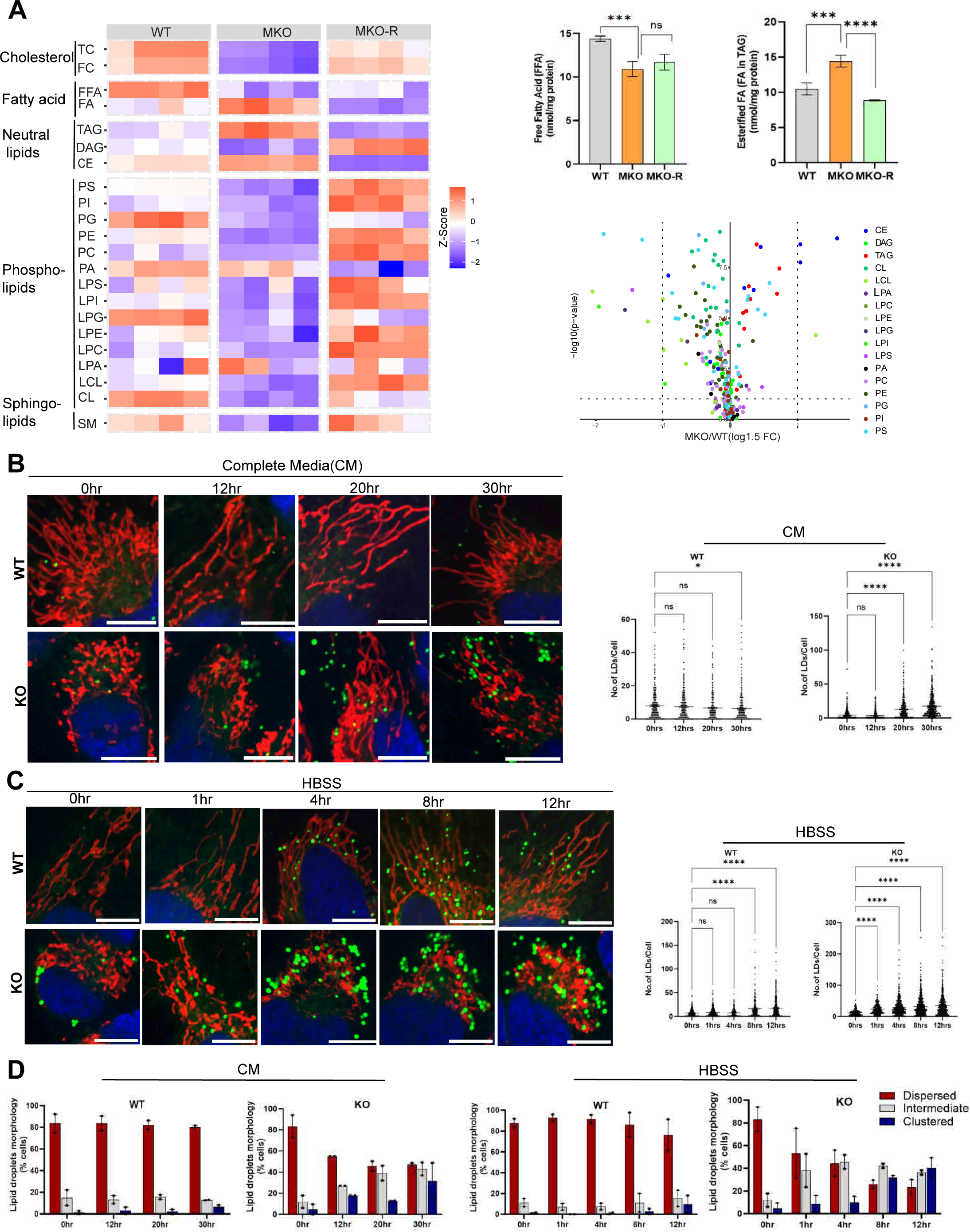
Membrane lipids decrease and storage lipids and lipid droplets increase in MKO cells. A. **Left panel:** Heat map comparing the levels of lipids at 20 hrs post media change in all 3 cell lines. Membrane lipids: Total Cholesterol (TC), free Cholesterol (FC), Phosphatidylserine (PS), Phosphatidylinositol (PI), Phosphatidylglycerol (PG), Phosphatidylethanolamine (PE), Phosphatidylcholine (PC), Phosphatidic acid (PA), Lyso-Phosphatidylserine (LPS), Lyso-Phosphatidylinositol (LPI), Lyso-Phosphatidylglycerol (LPG), Lyso-Phosphatidylethanolamine (LPE), Lyso-Phosphatidylcholine (LPC), Lyso-Phosphatidic acid (LPA), Lyso-Cardiolipin (LCL), Cardiolipin (CL), Sphingomyelin (SM). Neutral/Storage lipids: Triglycerides (TAG), Cholesterol ester (CE), Diglycerides (DAG). Fatty acids: esterified fatty acid (FA), free fatty acids (FFA). Cholesterol: Total Cholesterol (TC), Free Cholesterol (FC). We calculated the average value of all species per category per biological replicate. These values were compared between groups using ANOVA. Values in the heat map are scaled to z-scores per row (compound category). **Right top panel:** Average levels of esterified FA, FFA, in all 3 cell lines. Results in graphs are presented as mean ± SEM (\*\*\**p*<0.0003, \*\*\*\**p*<0.0007; One-way ANOVA, n=4 biological replicates). **Right bottom panel:** A Volcano plot comparing the levels of membrane and storage lipids between the WT and MKO cell lines. B. WT and MTCH2 KO HeLa cells were plated into complete media (CM), left to grow overnight, then media was changed (considered as time 0) and pictures were taken at 0, 12, 20, and 30 hrs post media change (left panel). LDs were labeled using BODIPY 493/503, and mitochondria were labeled using Mito Tracker deep red (MTDR). Right panel: Temporal quantification of the number of LDs in WT and MTCH2 KO cells at the four time points. Results are presented as means ± SEM (\**p*<0.05, \*\*\*\**p*<0.0007; n=3 biological replicates). C. WT and MTCH2 KO HeLa cells were plated into complete media, left to grow overnight, then media was changed to HBSS (considered as time 0) and pictures were taken at 0, 1, 4, 8, and 12 hrs post media change. Right panel: Temporal quantification of the number of LDs in WT and MTCH2 KO cells at the five time points. Data are presented as means ± SEM (\*\*\*\**p*<0.0007; n=3 biological replicates). D. Temporal quantification of the percentage of cells with dispersed, intermediate, or clustered LDs after incubation of cells for the indicated times in either: CM (left panel; pictures of cells appear in B) or HBSS (right panel; pictures of cells appear in C). Results are presented as means ± SEM. Scale bar=10μm.

**Table 2.** Results of ANOVA per compound group, followed by Tukey’s post-hoc test for pairwise comparisons. Each contrast is described by log-fold change, p-value and fdr-corrected p-value(Fig 3A).

TAGs and CEs are the major components of lipid droplets (LDs). LDs are an “on demand” energy source for the cell and can be mobilized in response to fluctuations in nutrient abundance^57^. In extended phases of nutrient scarcity, cells activate comprehensive strategies to collectively modify their metabolic processes, transitioning from predominantly using glycolysis to breaking down fatty acids through mitochondrial β-oxidation to produce energy. In accordance with the decrease in intracellular nutrients and membrane lipids, and the increase in TAGs and CEs, we found an increase in LD numbers and size in the MTCH2 knockout cells grown in complete media (Fig. 3B and Fig. S3C, respectively), which was further pronounced when cells were grown in HBSS nutrient depletion conditions (Fig. 3C). Notably, the pronounced accumulation of LDs in the MTCH2 knockout cells was accompanied by rearrangement of LDs from dispersed to a highly clustered distribution that was often observed in close proximity to mitochondria (Fig. 3D). Notably, we also found that MTCH2 knockout cells showed accelerated mitochondria elongation (Fig. S3D, top panels), which was further pronounced when cells were grown in HBSS (Fig. S3D, bottom panels).

Taken together, the accumulation of LDs in proximity to mitochondria and accelerated mitochondria elongation is likely to enable accelerated transfer and more efficient metabolism of lipid moieties at mitochondria resulting in increased energy production.

### MTCH2 is critical for adipocyte differentiation

Previously, it was reported that MTCH2 mRNA and protein expression are increased in obese women and during adipocyte differentiation^58^. It was also reported that deletion of MTCH2 inhibits adipogenesis and lipid accumulation in adipocyte cells^42,59^. Thus, adipogenesis/ lipogenesis is likely to be a physiologically relevant pathway to test the importance of MTCH2.

Adipose tissue plays a crucial role in regulating energy balance and glucose levels^60^, and the process of forming functional adipocytes involves the differentiation of preadipocytes into mature adipocytes. Moreover, NAD^+^ biosynthesis effectively integrates cellular metabolism with the adipogenic transcription program^25^. We used the CRISPR-Cas9 system to generate MTCH2 knockout NIH3T3L1 cells, a professional model to study adipogenesis. As expected, MTCH2 knockout in NIH3T3L1 cells resulted in mitochondrial fragmentation (Fig. S4A). During the process of differentiation in NIH3T3L1 preadipocytes, the WT proficient cells showed 80%-90% differentiation, based on lipid droplet quantification, whereas the MTCH2 knockout proficient cells showed 5%-10% differentiation on Day 6 (Fig. 4A).

**Figure 4.**
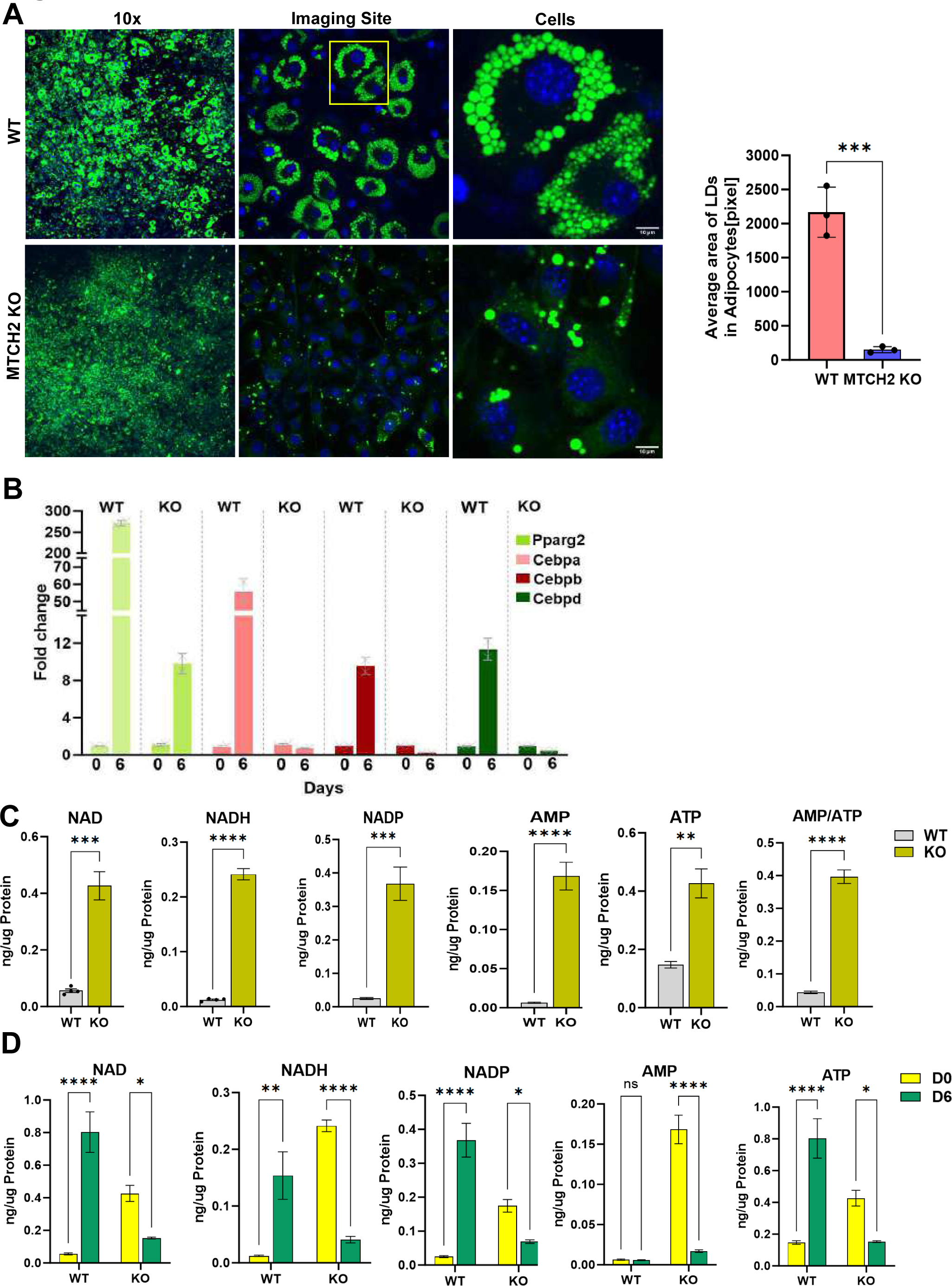
MTCH2 is critical for adipocyte differentiation. A. NIH3T3L1 cells were differentiated into adipocytes for 6 days in 4-well glass bottom plates. LDs were stained with Bodipy green and nuclei with Hoechst. The well overview is taken at 10X magnification. One region (marked by a yellow box) was magnified. Right panel: measure of differentiation by quantification of the number of LDs. Results are presented as mean ± SEM (\*\*\**p*<0.0003; One-way ANOVA, n=3 biological replicates). B. RT-PCR of WT and MTCH2 knockout (KO) cells at day 0 and day 6-post differentiation. Components of the adipogenic program Pparg, Cebpa, Cebpb, and Cebpd. Results are presented as mean ± SEM of one representative out of three independent experiments. Normalization was done by taking geometric mean of three housekeeping genes, Importin, Tubulin and AcTH. C. Levels of NAD^+^, NADH^+^, NADP^+^, AMP, ATP (and AMP/ATP ratio) in undifferentiated WT and MTCH2 KO preadipocyte NIH3T3L1. Results are presented as mean ± SEM (\**p*<0.05, \*\**p*<0.001, \*\*\**p*<0.0003, \*\*\*\**p*<0.0007, unpaired t-test, n=4 biological replicates). D. Levels of NAD^+^, NADH^+^, NADP^+^, AMP, ATP in WT and MTCH2 KO preadipocyte at day 0 and day 6-post differentiation. Results are presented as mean ± SEM (\**p*<0.05, \*\**p*<0.001, \*\*\*\**p*<0.0007, Two-way ANOVA with sidak’s multiple comparision test, n=4 biological Replicates).

As expected from the droplet quantification, Day 6 post-differentiation there was a substantial upregulation of the adipogenic transcription factors and many of their effectors in WT cells, which did not occur in the MTCH2 knockout cells (Fig. 4B and Fig. 4SB, respectively). Targeted metabolic profiling of NAD^+^, NADP^+^, NADH, AMP and ATP in resting preadipocytes prior to differentiation showed increased levels in the MTCH2 knockout cells as compared to the WT cells (Fig. 4C). High levels of ATP and of reducing equivalents are necessary for anabolic processes like lipid synthesis^61^. The higher levels of NAD^+^ and NADP^+^ suggests that MTCH2 knockout preadipocytes have an oxidizing intracellular environment that is inhibitory to anabolism, and thus inhibitory to reductive lipid biosynthesis^61^. Moreover, although MTCH2 knockout preadipocytes have higher ATP levels than the WT preadipocytes, their AMP levels were even higher, resulting in an AMP/ATP ratio that is ∼8-fold higher in the MTCH2 knockout as compared to the WT preadipocytes (Fig. 4C).

These results suggest that the MTCH2 knockout preadipocytes face a cellular energy crisis that is similar to the one seen in the MTCH2 knockout HeLa cells presented earlier. Moreover, targeted metabolomics comparing days 0 and days 6, post-differentiation, showed as expected that the levels of NAD^+^, NADP^+^, and ATP are increased in WT cells for proper signaling and to sustain enhanced anabolism during differentiation, whereas these metabolites are decreased in MTCH2 knockout cells (Fig. 4D). Thus, loss of MTCH2 results in a decrease in anabolic processes, like fatty acid biosynthesis, which are essential for adipogenesis.

## Discussion

In this study, we focused on understating the role of MTCH2 in metabolism. It is well-established that MTCH2 is a regulator of apoptosis by acting as the mitochondrial receptor for pro-apoptotic BID^29^, however its roles in regulating mitochondrial fusion and metabolism are less understood. Notably, conditional knockout of MTCH2 in mouse skeletal muscle results in protection from high fat diet-induced obesity, and this protection is likely due to an increase in whole-body energy utilization^30^.

How does loss of MTCH2 increase energy utilization? From our present results, we understood that MTCH2 balances the flow of energy among different metabolic pathways according to the cellular demand. MTCH2 seems to be involved in regulating the activity of several different metabolic pathways (Fig. 5). Proper timing and sustainable use of metabolic intermediates according to the cellular need is an indispensable process to run the cellular growth and proliferation. Under normal growth conditions, MTCH2, stationed on the surface of mitochondria like an antenna, is likely to be involved in the mitochondrial information processing system (MIPS)^7^. Mitochondria are the “headquarters” of cellular metabolism, harboring numerous metabolic inputs and outputs. Thus, MTCH2 might act like a “relay station” by sensing and connecting between metabolic intermediates/pathways and dynamic changes in mitochondria morphology/energy production by receiving and sending Wi-Fi signals (Fig. 5, left panel).

**Figure 5.**
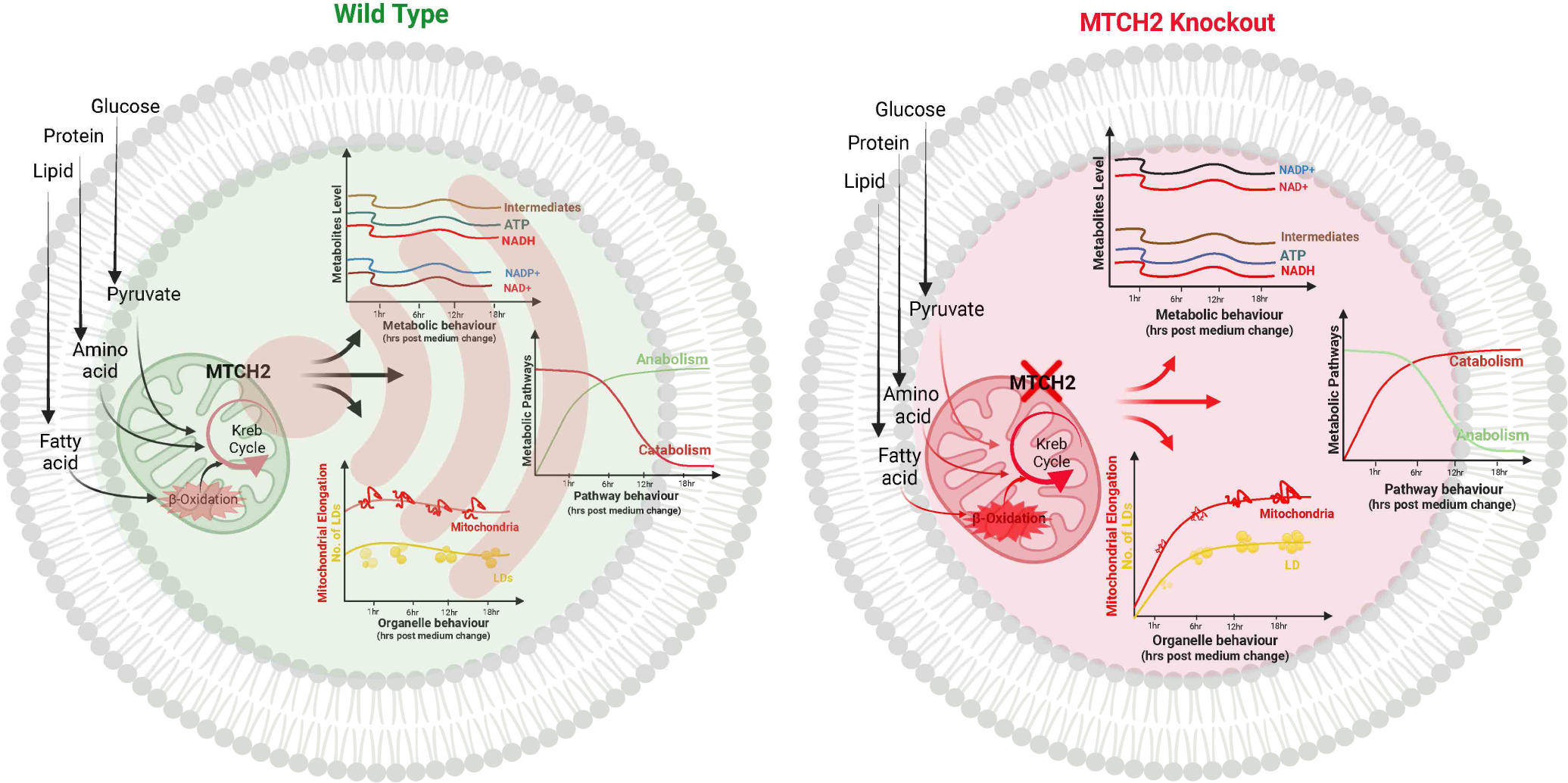
Schematic representation comparing the metabolic state of wild type and MTCH2 knockout cells. In wild type cells (left panel), MTCH2 might act as a mitochondrial “relay station” by sensing and connecting between metabolic intermediates/pathways and dynamic changes in mitochondria morphology/energy production by receiving and sending Wi-Fi signals. MTCH2 knockout, mimicking a scenario of losing a pivotal “relay station”, can lead to a disconnection between the cellular energy demand and the cellular energy utilization (created with BioRender.com).

The metabolomics analysis revealed that MTCH2 knockout results in an imbalance in several metabolic parameters: 1) Imbalance in the redox cofactors, NAD^+^, NADH and NADP^+^, leading to an oxidative environment. NAD^+^ plays an indispensable role in OXPHOS by acting as a proton acceptor^62^. Along with its role in OXPHOS it also acts as a signaling molecule in many cellular pathways like cell growth^63^, sirtuin activity^64^, and cell differentiation^24,25,65^. There is an intimate connection between energy metabolism and redox cofactors^12^, and they appear on the front defense line in incidents of mitochondrial insult^66^, cellular stress^67^, nutrient depletion/starvation^12^, and high energy demand like during exercise^68^. 2) Imbalance in adenine nucleotides, which results in an increase in the ADP/ATP ratio, representing an increase in energy demand. 3) Imbalance/decrease in many metabolites (carbohydrates, lipids, proteins, amino acids) and an increase in glucose uptake pointing to stimulation of metabolism/catabolism to meet the increased energy demands. Absence of MTCH2, mimicking a scenario of losing a pivotal “relay station” can lead to a disconnection between the cellular energy demand and the cellular energy utilization (Fig. 5, right panel).

Energy metabolism, redox potential and sustainable use of available nutrients dictates the “fingerprint” of cellular behavior^63,69^. Lack of coordination between the three systems will result in improper cellular growth and development^68^, and this might be the reason why MTCH2 knockout in mice results in embryonic lethality at E7.5^29^. In culture, MTCH2 knockout cells may survive at the cost of being smaller in size and growing slower^37^. The uncoordinated growth may also explain the delayed transition from naïve-to-prime in MTCH2 knockout embryonic stem cells and the increased exit from quiescence of MTCH2 knockout hematopoietic stem cells^31,37^. In culture, MTCH2 knockout cells seem to adapt by metabolic rewiring, which includes lipid rerouting to storage lipids to energize the cells instead of making membrane lipids for cell growth and proliferation.

As described above, oxidative metabolism is not an innate behavior but rather an adaptation of cells during specific conditions of cellular stress like starvation^70,71^. Since oxidative metabolism is not a canonical scenario and MTCH2 knockout leads to an oxidative environment, it is unfavorable to reductive biosynthesis pathways, like lipid synthesis^61^. Reductive biosynthesis pathways require a surplus amount of ATP and of the reducing cofactors NADH and NADPH to proceed^61,67^. NIH3T3L1 preadipocytes, a model to study white adipocyte physiology, revealed that MTCH2 knockout preadipocytes fail to differentiate into mature adipocytes. There can be at least three possible reasons to explain this phenotype: 1) mitochondrial fragmentation, 2) oxidative environment (increased levels of NAD^+^ and NADP^+^), and 3) shortage of metabolic intermediates.

Both the dynamic transition of mitochondria from a fragmented to a tubular state, and the metabolic transition from glycolysis to OXPHOS are important to meet the increased energy demand for anabolic processes during differentiation^24,65^. To make one molecule of fatty acid, palmitic acid, cells need approximately 14 molecules of NADPH and 7 molecules of ATP along with 16 carbons from 8 molecules of acetyl CoA^72^. Targeted metabolomics of the MTCH2 knockout NIH3T3L1 preadipocytes showed higher levels of NAD^+^, NADP^+^, and a higher AMP/ATP ratio, indicating an oxidative and low-energy environment, which is not favorable for differentiation. In addition, the nuclear pool of NAD^+^ acts as a signaling molecule and an increase in the cytoplasmic NAD^+^ levels are used to maintain metabolic intermediates of glucose metabolism during differentiation^25,65^. Our results show that during differentiation (day 0 to day 6) WT cells increase, whereas the MTCH2 knockout cells decrease, their NAD^+^ levels.

How does MTCH2 balance the energy flow in cells? It was recently reported that MTCH2 regulates mitochondrial fusion by modulating the pro-mitochondrial fusion lipid lysophosphatidic acid (LPA)^38^. Most recently, we found that MTCH2 cooperates with ER-localized MFN2 and LPA synthesis at the ER to sustain mitochondrial fusion (Goldman et al, *EMBO Rep* in Press). Thus, MTCH2 may play a role in phospholipid transfer from the ER to mitochondria. MTCH2 was also demonstrated to act as an insertase, which aids tail-anchored mitochondrial proteins to integrate into the OMM^43^. Interestingly, mutations in the central predicated “pore” region of MTCH2 led to either an increase or a decrease in its insertase activity^43^. It was also reported that insertases can function as scramblases (including MTCH2), which flip phospholipids between the two leaflets of the membrane through their “pore” region^44,45^. Thus, MTCH2 may balance cellular energy flow by regulating the mitochondria membrane lipid composition. Moreover, this putative lipid-modifying activity might be also related to MTCH2’s apoptotic activity in regulating tBID-induced cytochrome c release from mitochondria.

In summary, we show that knockout of MTCH2 results in an unbalanced energy flow in cells. Loss of MTCH2 stimulates many metabolic pathways to meet the unbalanced cellular demand for ATP. These findings are consistent with the idea that MTCH2 is a critical regulator of energy flow in cells.

### Lead contacts and materials availability

Further information and requests for resources and reagents should be directed to and will be fulfilled by the lead contacts Atan Gross and Sabita Chourasia.

### Experimental model and subject details

#### Cell lines

##### HeLa Cells

HeLa cells were cultured in Dulbecco’s Modified Eagle’s Medium (DMEM) containing 4.5 g/l glucose and L-glutamine (cat. # 41965, Gibco), supplemented with sodium pyruvate (cat. # 03-042, Biological Industries) and 10% fetal bovine serum (FBS; cat. # 12657, Gibco), at 37°C and 5% CO_2_. Complete media (CM) conditions consisted of DMEM with 4.5 g/l glucose and L-glutamine, supplemented with sodium pyruvate and 10% fetal bovine serum (FBS). Nutrient depletion conditions consisted of growth in Hank’s Balancing Salt Solution (HBSS; cat. # 02-015, Biological Industries).

##### NIH3T3L1 Cells

NIH3T3L1 Preadipocytes cells (ATCC) were cultured in Dulbecco’s Modified Eagle’s Medium (DMEM) containing 4.5 g/l glucose and L-glutamine (cat. # 41965, Gibco), supplemented with 10% fetal calf serum (FCS; cat. # 04-102-1A, Biological Industries) at 37°C and 5% CO_2_. The cells were cultured in 10 cm tissue culture dishes (TPP) and the medium was replaced every 2 days. The pre-adipocytes never reached densities above 70% confluence unless they were subjected to differentiation (described below).

### Preparation of adipocyte differentiation medium

The differentiation medium utilized in these experiments was comprised of Dulbecco’s Modified Eagle’s Medium (DMEM; cat. # 41965, Gibco) and 10% fetal bovine serum (FBS; cat. # 12657, Gibco). To induce differentiation, the medium was supplemented with specific reagents: 517 mmol/L 3-Isobutyl-1-methylxanthine (IBMX; Sigma-Aldrich), 1 mmol/L dexamethasone (G-biosciences), and 167 nmol/L bovine insulin (Sigma-Aldrich). For the preparation of the differentiation medium, a fresh 100-fold IBMX stock solution was created for each experiment at a concentration of 0.0115 g/mL in 0.5 M KOH (Merck). Bovine insulin was prepared from a 10,000-fold stock solution at a final concentration of 10 mg/mL, following the manufacturer’s specifications (Sigma-Aldrich). Additionally, a 1,000-fold dexamethasone stock solution was prepared by diluting tenfold the provided 10 mM solution from the manufacturer (G-biosciences) with phosphate-buffered saline solution (PBS; Thermo-Fisher). Before use, the IBMX stock solution was diluted 1:100, the insulin stock solution was diluted 1:10,000, and the dexamethasone stock solution was diluted 1:1,000 in the appropriate volume of DMEM with 10% FBS. All solutions underwent sterile filtration to ensure aseptic conditions throughout the experiments.

### Adipocyte differentiation in 4-well glass bottom imaging plate

NIH3T3L1 pre-adipocytes were initially seeded at a density of 10,000 cells per well in 4-well glass bottom imaging plates and cultured for 2 days until they achieved complete confluence. The culture medium was then replaced with fresh DMEM supplemented with 10% FCS. After 48 hours, the cells underwent differentiation using the previously detailed differentiation medium. Six days into the differentiation process, the cells were stained with BODIPY 493/503 (cat. # D3922, Thermo Scientific) and Mito Tracker Deep Red (MTDR; cat. # M22426, Invitrogen) before being processed for imaging.

### Adipocyte differentiation in 6 well format

NIH3T3L1 pre-adipocytes were seeded at a density of 200,000 cells per well in 6-well plates and cultured for 2 days until they achieved complete confluence. The culture medium was then replaced with fresh DMEM supplemented with 10% FCS. After 48 hours, the cells underwent differentiation using the same protocol employed for 10 cm tissue culture dishes, as described below.

### Adipocyte differentiation in 10 cm tissue culture dishes

The cells were maintained following the procedure. To initiate differentiation, cells were cultured until they reached complete confluence and then further incubated for an additional 48 hours. Once the cells achieved 100% confluence after 48 hours, the medium was replaced with the differentiation medium, prepared as previously described. This time point, marking the addition of the differentiation medium, was defined as the start of differentiation (day 0). After two days, the differentiation medium was substituted with DMEM/10% FBS containing 167 nmol/L insulin. On day 4 post-initiation of differentiation, the medium was replaced with DMEM/10% FBS. Finally, at day 6 post-initiation of differentiation, the cells were harvested.

### Generation of MTCH2 knockout stable cell lines

#### Generation of MTCH2 knockout cells using the CRISPR Cas9 lentiviral system

The MTCH2 CRISPR KO cell line was generated in HeLa and NIH3T3L1 cells. Guides were designed using CHOP web application (https://chopchop.cbu.uib.no/). The MTCH2 guides-RNAs were F-CACCAGCACTTTCACGTACATGAGGT and R-TAAAACCTCATGTACGTGAAAGTGCT. gRNAs were cloned in pKLV-U6gRNA (BbsI)-PGKpuro2ABFP (Plasmid #50946). To generate CRISPR-Cas9 knockout cell lines, HeLa and NIH3T3L1 cells were co-transfected with 1) gRNA containing plasmid and 2) GFP-Cas9 (pCas9_GFP, Plasmid #44719). To control for effects of CRISPR-Cas9 expression on HeLa and NIH3T3L1 cells, we generated a CRISPR-Cas9 control cell line expressing the same CRISPR-Cas9 construct, but without gRNA. Positively transduced cells were selected in DMEM, 10% FBS and 2μg/ml puromycin for 2 weeks.

#### Generation of MKO-R cells by expressing MTCH2 in MKO cells using a retroviral system

To generate HeLa cells carrying a stable MTCH2 gene, we used a pBaBe based retroviral construct for gene transfer. MTCH2 was sub-cloned into the ENTRY vector (pBaBe; Addgene) using TOPO cloning. Stable cell pools were generated by selecting positively transduced cells with puromycin selection (2μg/ml) for two weeks.

#### Production of retroviruses for gene transfer

The retrovirus for gene expression was generated in HEK293T cells. HEK293T cells, maintained in DMEM with 10% FBS and supplemented with sodium pyruvate, were cultured in 6 cm dishes until reaching 70% confluence. Transfection of HEK293T cells was carried out using jet-PI. Briefly, cells were co-transfected for 6 hours with three plasmids: 10 μg of a plasmid containing the expression constructs, 6.5 μg of the viral genome packaging plasmid psPAX2, and 3.5 μg of pMD2.G, a plasmid producing the virus envelope components. The plasmids were mixed in double-distilled water (ddH2O) to a final volume of 597 μl. After 30 minutes at room temperature, the transfection mix was added to the cells. The media was changed after 12 hrs and replaced with 10 mL of DMEM with 10% FBS. Media was collected and replaced 24 and 48 hrs after transfection. Each fraction was filtered through a 0.45 μm sterile filter and stored at 4°C until use. To concentrate the virus, all fractions were combined and centrifuged using an ultrafiltration centricon (Amicon "Ultra – 15 centrifugal filter" with a 100 kDa MW cutoff) for 45 minutes at 4°C. The filtrate was resuspended in a total volume of 1 mL (DMEM, 10% FBS), which was then used to infect the HeLa MTCH2 KO cells.

#### Plasmid transfection of HeLa and NIH3T3L1 cells

For transfection, HeLa and NIH3T3L1 cells were cultured in 6-well plates with DMEM, 10% FBS to 80% confluence. The cells were then transfected using the transfection reagent Lipofectamine 3000 (Thermo-Fisher) following the instructions of the manufacturer. Transfection reagent and DNA were prediluted in Opti-MEM medium (Cat. #31985062, Gibco). The DNA transfection reagent complex was allowed to form for 15 min at room temperature and then transferred dropwise into the culture medium. The cells were incubated in the presence of the transfection mix for another 6 hrs at 37°C in DMEM/10% FBS and then media was replaced by fresh media containing DMEM/10% FBS and on the next day media was replaced by media containing puromycin (2μg/ml) followed by 2 weeks of selection.

#### Quantification of the adipocyte differentiation efficiency

Image Processing, analysis, and statistics images were analyzed using ImageJ (NIH). LD size and number were quantified with the ImageJ ‘‘analyze particles’’ function in thresholded images, with size (square pixel) settings from 0.1 to 100 and circularity from 0 to 1. Average area of LDs was analyzed. Data were expressed as means ± SEM. Statistical analysis among groups was performed using Student’s t test.

#### Quantitative real-time PCR

Samples were collected on Day0 and Day7 of differentiation. RNA was extracted using NucleoSpin RNA kit (Macherey-Nagel #740955) following the manufacturer’s instructions. A sample corresponding to 1μg RNA from each sample was used to perform cDNA synthesis by the High-Capacity cDNA Reverse Transcription Kit (Cat. # 4368814, Applied Biosystems). qPCR was performed using 0.4 ng/μl cDNA and 0.5 μM of each primer, whose sequences are listed in Table 3.

**Table 3.** List of primers used in Fig4 and FigS4.

### Fluorescence Microscopy

#### Live imaging

For live cell imaging experiments, Hela cells were seeded a day before the experiment. Media was replaced by either fresh Complete Media (CM) or HBSS and considered as time=0. Cells were pre-incubated with 100 nM Mito tracker Deep Red (MTDR; cat. # M22426) for mitochondria staining and with 1μg BODIPY 493/503 for LD staining (for 30 min). Since, our aim was to see changes over time, we maintained the same media until the imaging was complete. After staining, cells were stabilized for an additional 30 min at 37°C and 5% CO_2_. Cells were then imaged under temperature and CO_2_-controlled conditions. Stained cells were analyzed using a Nikon ECLIPSE Ti2-E inverted microscope with a CSW-1 spinning disc system (Yokogawa), and with a x100 CFI Plan Apo100x oil (na 1.45 wd 0.13mm), equipped with temperature and CO_2_ control. Cells were incubated at 37°C in a 5% CO_2_ humidified chamber and images were taken at 0, 1, 12, 20, 30 hrs for CM and 0, 1, 4, 8, 12 hrs for HBSS.

#### Automated image analysis of live imaging experiments with HeLa cells

All images were analyzed with open-source soft wares and we used Fiji^73^, StarDist^74^, Ilastik^75^ and Cellpose^76^. Below we describe the main steps taken.

##### 1) Single Cell segmentation by Cellpose

To identify individual cells in the image we trained a Cellpose model using both the mitochondria and dapi channels. The training was done on representative images of different conditions, from both WT and MTCH2 KO groups. We then dilated the identified cells as to include the cell’s membrane.

##### 2) LD segmentation and clustering by StarDist and Cellpose

To identify LDs, we used StarDist for the MTCH2 KO group. For the WT group, we used StarDist or Cellpose alternatively since at certain time points (early hours of post-media change), the LDs have low intensity and StarDist fails to identify them. Cellpose segmentation is better at identifying the LDs at these time points, but still has high false positives. To avoid these false positives, we filtered LDs based on their mean intensity (keeping only the top 10-20%). For quantitating LD clustering, we used Fiji’s SSIDC cluster indicator plugin.

##### 3) Pixel based mitochondria segmentation by Ilastik

To segment mitochondria, we trained an Ilastik model using representative images of all different conditions from both WT and MTCH2 KO groups.

##### 4) Cell categorization

For each cell identified in each image in, we exported the related LD and Mitochondria information to an Excel spreadsheet. We used the excel spreadsheet to categorize cells according to the size of the mitochondria length and level of LD clustering.

### LC-MS based targeted metabolomics

Frozen cell lysates were aliquoted and extracted in organic extraction solvents for targeted LC/MS metabolomics (acylcarnitines, amino acids, organic acids, nucleotides, and malonyl and acetyl CoA) according to validated, optimized protocols in our previously published studies^77,78^. These protocols use cold conditions and solvents to arrest cellular metabolism and maximize the stability and extraction recovery of metabolites. Each class of metabolites was separated with a unique HPLC method to optimize their chromatographic resolution and sensitivity. Quantitation of metabolites in each assay module was achieved using multiple reaction monitoring of calibration solutions and study samples on an Agilent 1290 Infinity UHPLC/6495 triple quadrupole mass spectrometer^77,78^. Raw data was processed using Mass Hunter quantitative analysis software (Agilent). Calibration curves (R2 = 0.99 or greater) are either fitted with a linear or a quadratic curve with a 1/X or 1/X2 weighting.

### Non-Targeted (Global) metabolomics

#### Metabolite extraction

Extraction and analysis of polar metabolites were performed as previously described^79,80^ with a few modifications: Samples were lyophilized and extracted with 1ml of a pre-cooled (−20°C) homogenous methanol:methyl-tert-butyl-ether (MTBE) (1:3, v/v) mixture. The tubes were vortexed and then sonicated for 30 min in an ice-cold sonication bath (taken for a brief vortex every 10 min). Then, DDW:methanol (3:1, v/v) solution (0.5ml), containing internal following standards: C13 and N15 labeled amino acids standard mix (Sigma, 767964) (1:500), was added to the tubes followed by vortex and centrifugation. The upper organic phase was removed, and the lower polar phase was re-extracted as described above, with 0.5ml of MTBE, moved to a new Eppendorf tube, dried in speed vac, and stored at −80°C until analysis. For analysis, the polar dry samples were re-suspended in 150µl methanol:DDW (50:50) and centrifuged twice to remove the debris. 125µl were transferred to the HPLC vials for injection.

#### LC-MS polar metabolite analysis

Metabolic profiling of the polar phase was done as described^79^, with minor modifications. Briefly, analysis was performed using Acquity I class UPLC System combined with mass spectrometer Q Exactive Plus Orbitrap™ (Thermo Fisher Scientific) operated in a negative ionization mode. The LC separation was done using the SeQuant Zic-pHilic (150 mm × 2.1 mm) with the SeQuant guard column (20 mm × 2.1 mm) (Merck). The Mobile Phase B: acetonitrile and Mobile Phase A: 20 mM ammonium carbonate with 0.1% ammonia hydroxide in water:acetonitrile (80:20, v/v). The flow rate was kept at 200μl* min^−1,^ and the gradient was as follows: 0-2min 75% of B, 14 min 25% of B, 18 min 25% of B, 19 min 75% of B, for 4 min, 23 min 75% of B.

#### Polar metabolites data analysis

The data was processed using Progenesis QI (Waters) when detected compounds were identified by accurate mass, retention time, isotope pattern, and fragments and verified using an in-house-generated mass spectra library.

#### Shotgun lipidomics

Lipid species were analyzed using multidimensional mass spectrometry-based shotgun lipidomic analysis^81^. In brief, each cell sample homogenate containing 0.5mg of protein, which was determined with a Pierce BCA assay was accurately transferred to a disposable glass culture test tube. A premixture of lipid internal standards (IS) was added prior to conducting lipid extraction for quantification of the targeted lipid species. Lipid extraction was performed using a modified Bligh and Dyer procedure^81^, and each lipid extract was reconstituted in chloroform:methanol (1:1, *v:v*) at a volume of 400µl/mg protein. Phosphoethanolamine (PE), cholesterol (CHL), free fatty acid (FFA) and diacylglycerol (DAG) were derivatized as described previously^82–85^ before lipidomic analysis. Lysophospholipids (LPA, LPG, LPI and LPS) in water phase were enriched using HybridSPE cartridge, after washing with methanol, the lysophospholipids were eluted with methanol/ammonia hydroxide (9:1 and 8:2), dried and reconstituted in methanol for lipidomic analysis^85^.

For shotgun lipidomics, lipid extract was further diluted to a final concentration of ∼500 fmol total lipids per µl. Mass spectrometric analysis was performed on a triple quadrupole mass spectrometer (TSQ Altis, Thermo Fisher Scientific, San Jose, CA) and a Q Exactive mass spectrometer (Thermo Scientific, San Jose, CA), both of which were equipped with an automated nanospray device (TriVersa NanoMate, Advion Bioscience Ltd., Ithaca, NY) as described^86^. Identification and quantification of lipid species were performed using an automated software program^87,88^ Data processing (e.g., ion peak selection, baseline correction, data transfer, peak intensity comparison and quantitation) was performed as described^88^. The results were normalized to the protein content (nmol lipid/mg protein).

#### Targeted metabolomics for ATP, AMP, NAD, NADH, NADP in NIH3T3L1 cells Materials

Adenosine 5’-triphosphate (ATP), adenosine 5’-diphosphate (ADP), adenosine 5’-monophosphate (AMP), β-Nicotinamide adenine dinucleotide (NAD^+^), β-Nicotinamide adenine dinucleotide reduced (NADH), β-Nicotinamide adenine dinucleotide phosphate (NADP), ^13^C_10_-adenosine 5’-triphosphate (^13^C_10_-ATP), ^15^N_5_-adenosine 5’-monophosphate (^15^N_5_-AMP), and amino acid internal standard mix - all were purchased from Merck.

#### Sample preparation

Dried pellet of 20 million cells was extracted with 400μl of 10mM ammonium acetate and 5mM ammonium bicarbonate buffer, pH 7.7 and 600μl methanol in bead beater (10Hz, 1min; Retsch MM400) and next in shaker (1,200rpm, 30min; Thermomixer C Eppendorf). Then the extract was centrifuged (19,000g, 10min), the supernatant collected and evaporated under reduced pressure. The obtained residue was re-dissolved in 100μl of 50%-aqueous acetonitrile and placed in LC-MS filter vial (0.2-um PES, Thomson).

#### Liquid Chromatography - tandem Mass Spectrometry (LC-MS/MS)

LC-MS/MS analysis was performed using an instrument consisted of Acquity I-class UPLC system (Waters) and Xevo TQ-S triple quadrupole mass spectrometer (Waters).

**LC:** Metabolites were separated on an Atlantis Premier Z-HILIC column (2.1 × 150 mm, 1.7 μm particle size; Waters). Mobile phase consisted of (A) 20% acetonitrile in 20mM ammonium carbonate buffer, pH 9.25 and (B) acetonitrile. Gradient conditions were: 0 to 0.8 min = 80% B; then to 5.6 min gradient (curve 3) to 25% B; 5.6 to 6 min = hold at 25% B; 6 to 6.4 min = back to 80% B. Total run time 9 min. Injection volume was 3μl, and flow rate was 0.3ml/min.

**MS/MS:** Desolvation temperature 400°C, desolvation gas flow 800L/h, cone gas flow 150L/h nebulizer pressure 4 Bar, capillary voltage 2.49kV, collision gas (argon) flow 0.25 mL/min, source temperature 150°C. The MRM transitions used were: ATP: 507.9>136 m/z at collision 35V and cone voltage 14V; ADP: 428>136.05 and 428>348.1 m/z at collision 25 and 14V, respectively, and cone voltage 25V; AMP: 348>96.8, 348>119, and 348>135.7 m/z at collision 28, 55, and 20V, respectively, and cone voltage 10V; NAD: 664>136, 664>428 m/z at collision 37 and 27V, respectively, and cone voltage 10V; NADH: 666>514, 664>649 m/z at collision 30 and 20V, respectively, and cone voltage 10V; NADP: 744>508, 664>604 m/z at collision 28 and 20V, respectively, and cone voltage 10V; GSH: 308>179 m/z at collision 17V and cone voltage 10V; GSSG: 613>355 m/z at collision 32V and cone voltage 10V. Internal standards: ^13^C_10_-ATP: 518.12>141.07, 518.12>420.1 m/z at collision 35 and 18V, respectively, and cone voltage 14V; ^15^N_5_-AMP: 353.1>141.1 m/z at collision 20V and cone voltage 17V. MassLynx and TargetLynx software (v. 4.2 Waters) were applied for quantitative analysis using standard curve in 0.001–10 μg/ml concentration range for each metabolite. ^13^C_10_-ATP and ^15^N_5_-AMP were added to standards and samples as internal standards to get 0.1 and 0.5 μM, respectively.

#### Experimental Design

Experiments were not done in a blinded fashion. All instances where n replicates are reported had n biological replicates.

### Quantification and statistical analysis

#### Metabolomics and lipidomics

Comparisons of compounds or categories between the 3 groups were done using one-way ANOVA on log2-transformed values, followed by Tukey’s post-hoc test. All statistics were done in R, v. 4.3.1. Graphs were made in GraphPad Prism 10. Heat maps and Volcano plot were generated using ggplot2, v. 3.4.4. Metabolomics and Lipidomics analyses corrected p-value excel file appear in supplementary Tables 1 and 2, respectively. All instances where n replicates are reported had n biological replicates.

#### Fluorescence Microscopy

Fluorescence microscopy images were acquired using the Nikon ECLIPSE Ti2-E inverted microscope with CSW-1 spinning disc system (Yokogawa), with a x100 CFI Plan Apo100x oil (na 1.45 wd 0.13mm), equipped with temperature and CO_2_ control and were quantified using ImageJ software^73^. Images were thresholded, the area of BODIPY 493/503 stained LDs were quantified from three biological replicates (average of 50-100 cells per replicate), and the mean ± SEM was determined. Statistical significance was evaluated using the student t test with a p-value < 0.05. Data was presented as scattered plot generated in GraphPad Prism Software. Statistical significance was evaluated using the student t test with a p-value < 0.05.

#### RT-qPCR

The mean ± SEM was determined from three independent experiments. Statistical significance was evaluated using the student t-test with a p-value < 0.05.

#### Data and code availability

The Fiji macro used to run the analysis, and the Excel template used to categorize the individual cells, together with the trained Ilastik and Cellpose models are deposited and available for download on GitHub (link to be provided upon paper acceptance). Fiji can be downloaded from https://imagej.net/software/fiji/downloads. It includes ready to use StarDist and can be configured to run the BioVoxxel plugin that implements the SSCID clustering algorithm; Cellpose can be downloaded from https://github.com/mouseland/cellpose; and Ilastik can be downloaded from https://www.ilastik.org/download.

#### Author contributions

S. Chourasia performed all the experiments presented in the paper. S. Chourasia and A. Gross planned the experiments and wrote the manuscript. C. Petucci, A. Brandis, S. Malitsky, M. Itkin, and T. Mehlman performed the metabolomics, X. Han performed the lipidomics, E. Sivan and R. Rotkopf assisted with data analyses, and L. Regev and Y. Zaltsman assisted in performing some of the experiments. All authors discussed the results and commented on the manuscript.

## Acknowledgments

We are grateful to all the members of the Gross lab for their support, insightful discussions, and comments on the manuscript. Also, thank you to Dr. Nishanth Belugali Nataraj for his technical support during certain experiments and Dr. Reeba Jacob for discussion regarding image analysis.

---

**Figure S2.**
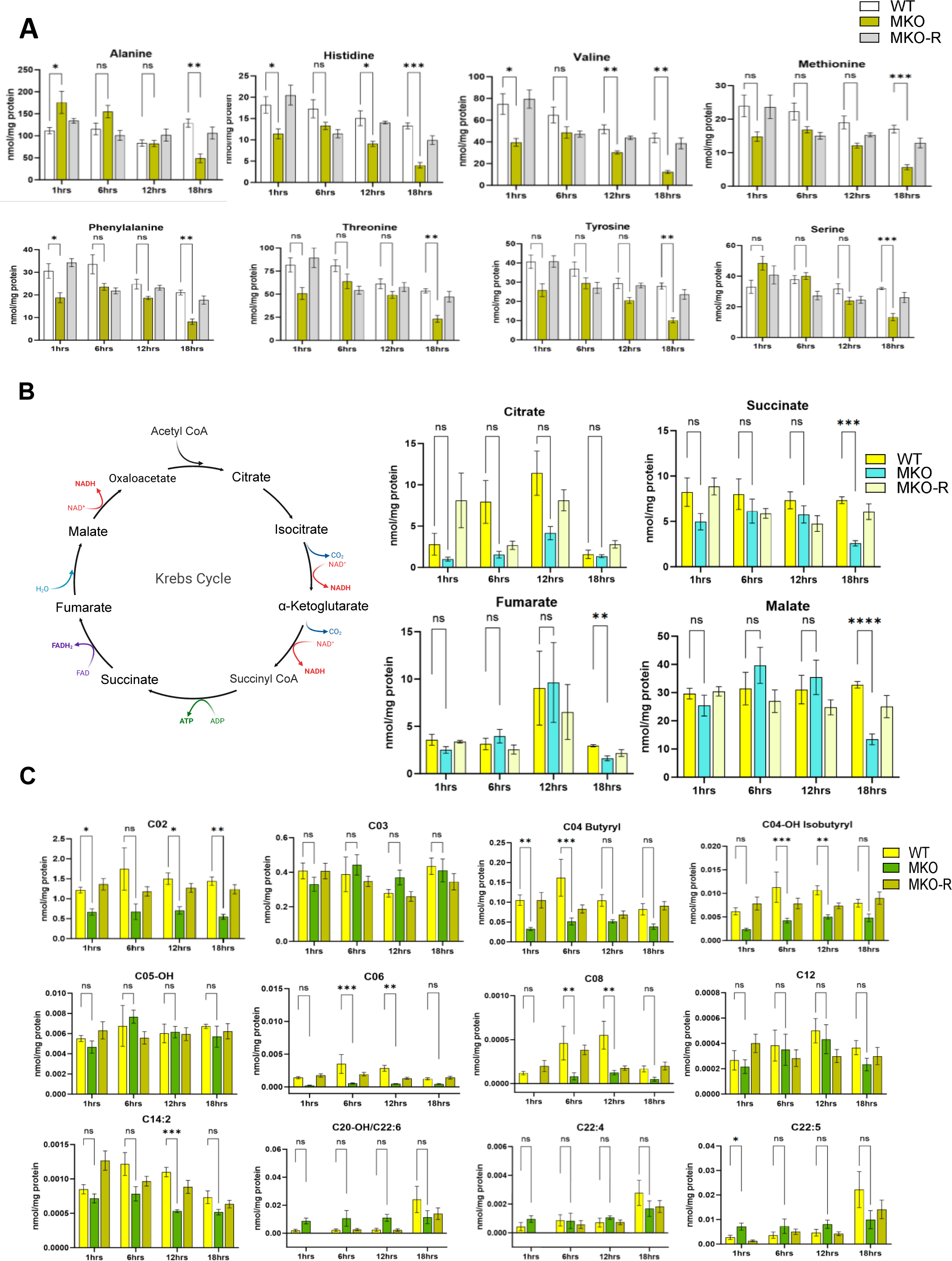
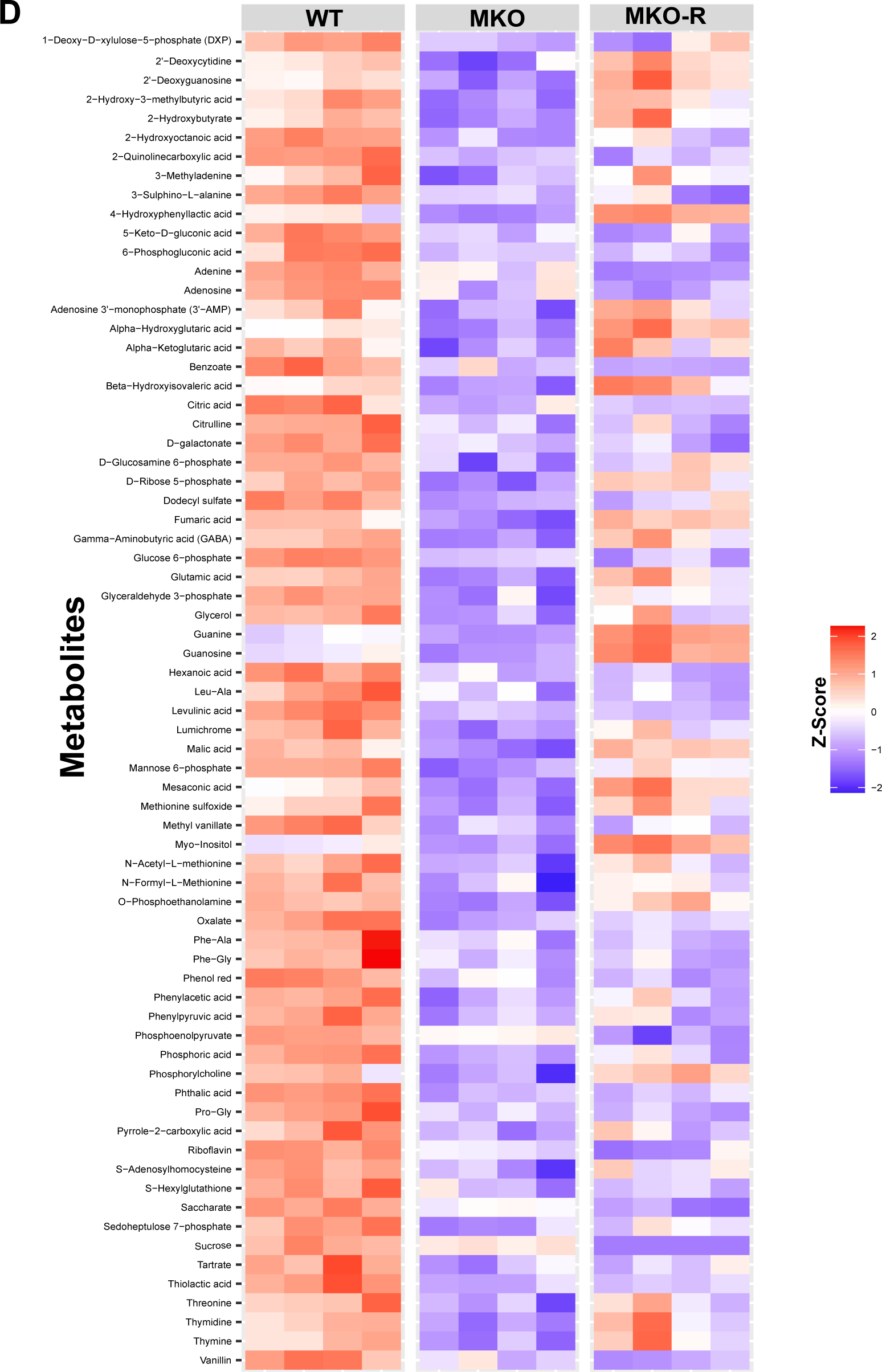
An increase in amino acid/TCA cycle/lipid metabolism and a decrease in many metabolites in MKO cells. A. Average levels of a set of amino acids in all 3 cell lines at all four time points. B. Left panel: Schematic representation of the TCA cycle. Right panels: Average levels of a set of TCA cycle intermediates in all 3 cell lines at all four time points. C. Average levels of acyl carnitines in all 3 cell lines at all four time points. Results in all graphs in A-C are presented as mean ± SEM (\**p*<0.05, \*\**p*<0.001, \*\*\**p*<0.0003, \*\*\*\**p*<0.0007; two-way ANOVA with Dunnett multiple comparison test; n=6 biological replicates). D. A heat map comparing the levels of 70 metabolites in WT, MKO and MKO-R cell lines. Metabolite concentration values (Relative abundance) were log1.5-transformed for statistics. The groups were compared by ANOVA. Values are scaled to z-scores per row (metabolite); n=4 Biological Replicates.

**Figure S3.**
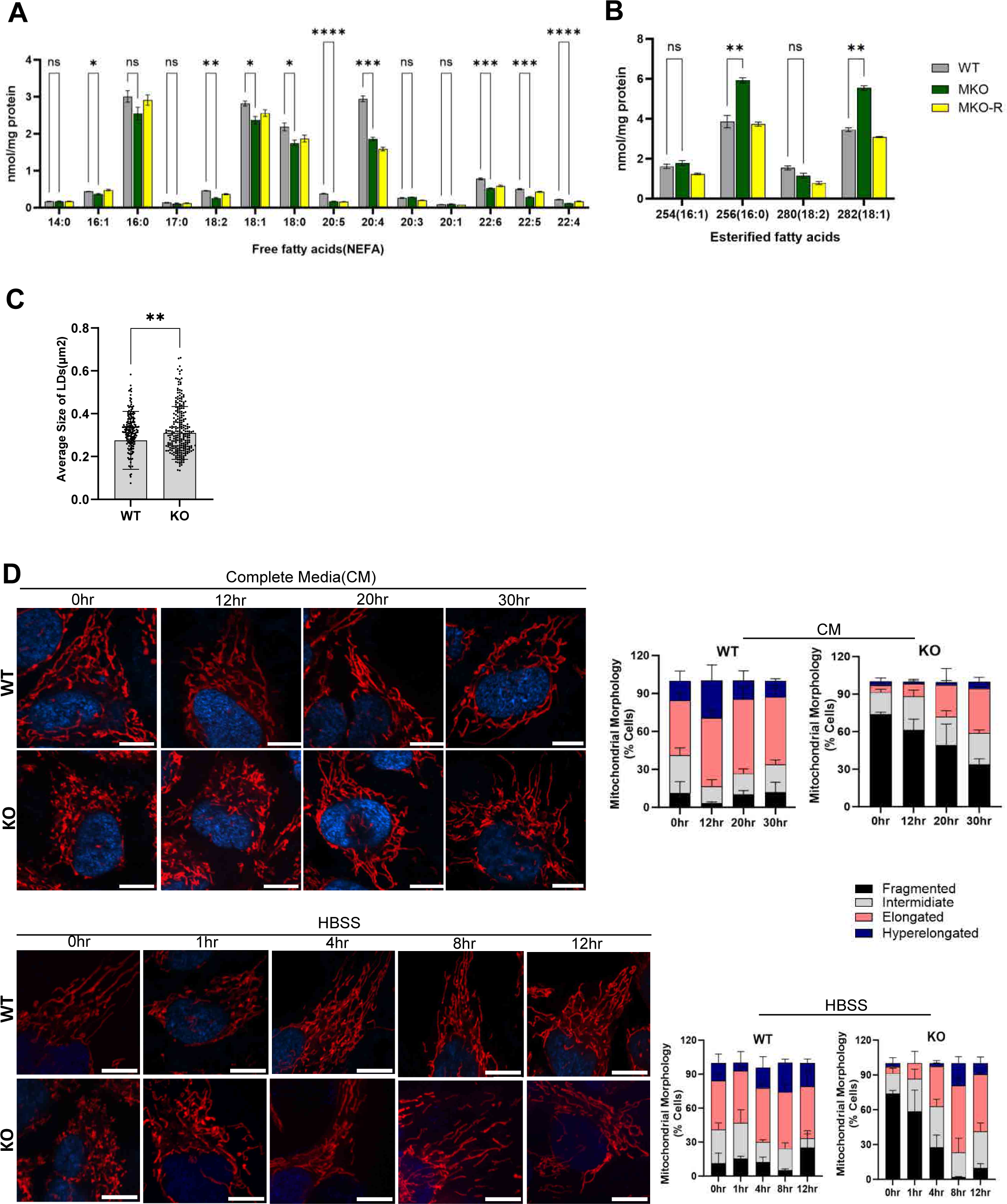
MTCH2 knockout cells show accelerated mitochondria elongation under nutrient depletion conditions. A, B. The levels of Free fatty acids (NEFA)(A) and esterified fatty acids (B) in all 3 cell lines. Results are presented as mean ± SEM (\**p*<0.05, \*\**p,* <0.001, \*\*\**p*<0.0003; One-way ANOVA, n=4 biological replicates). C. Quantification of LD average size in WT and MTCH2 KO cells. Results are presented as mean ± SEM (\*\**p,* <0.001; One-way ANOVA, n=3 biological replicates). D. Analyses of mitochondria morphology. WT and MTCH2 KO HeLa cells were plated into complete media (CM), left to grow overnight, then media was changed to CM (top) or HBSS (bottom)(considered as time 0) and pictures were taken at 0, 12, 20, and 30 hrs post media change. Mitochondria were labeled using Mito Tracker Deep red (MTDR). Right panels: Mitochondria morphology quantification of cells incubated in either CM (top) or in HBSS (bottom). Results are presented as means ± SEM (n=3 biological replicates). Scale bar=10μm.

**Figure S4.**
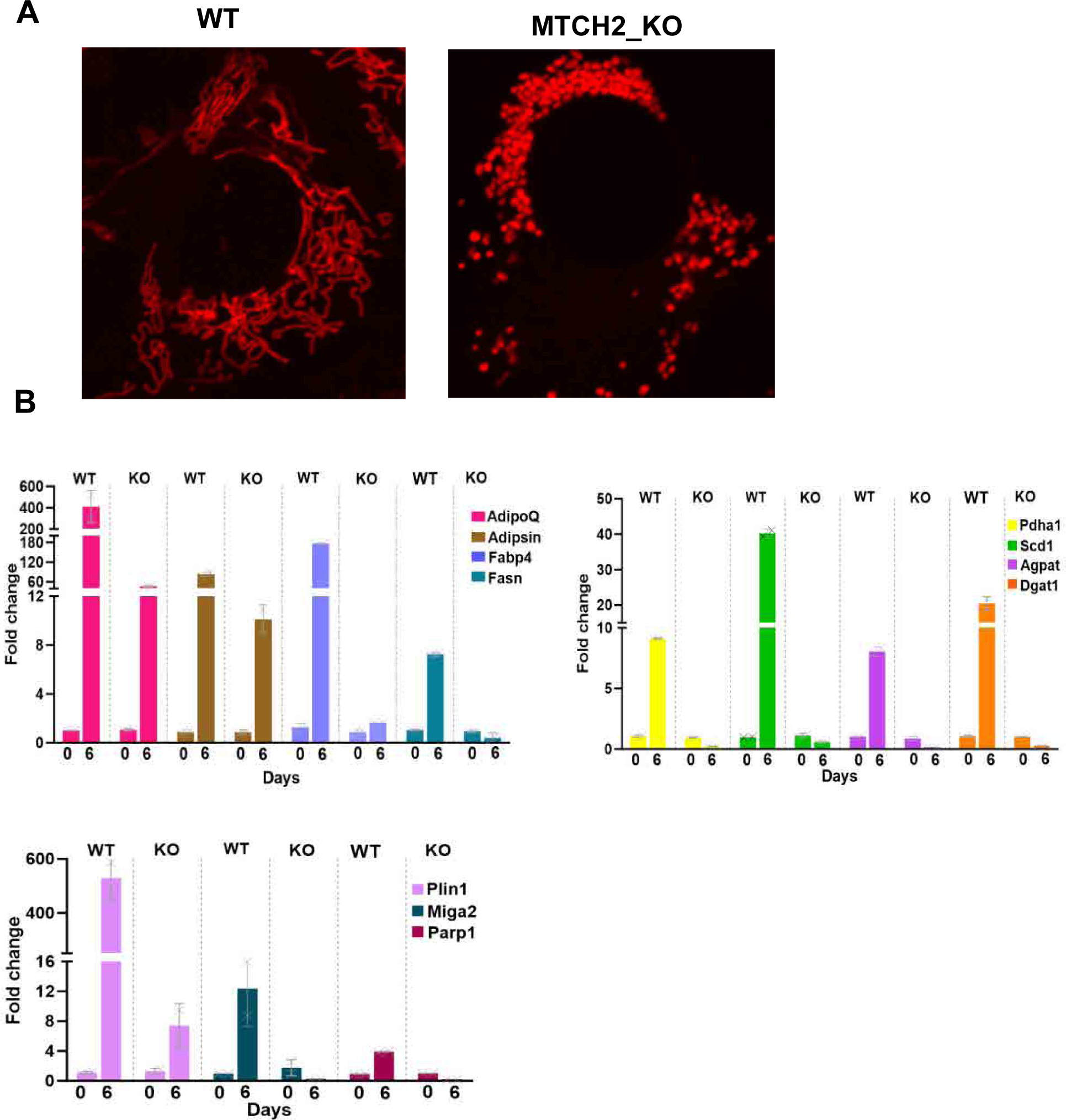
MTCH2 is critical for adipocyte differentiation. A. Mitochondrial morphology in NIH3T3L1 Preadipocytes. MTCH2 knockout (KO) leads to mitochondrial fragmentation (right panel). B. RT-PCR of WT and MTCH2 knockout (KO) cells at day 0 and day 6-post differentiation. Components of the adipogenic effector genes were analyzed: adiponectin (AdipoQ), Adipsin, fatty-acid-binding protein 4 (Fabp4), fatty acid synthase (FASN), pyruvate dehydrogenase (Pdha1), stearyl-CoA desaturase (Scd1), 1-acyl-sn-glycerol-3-phosphate (Agpat), diacylglycerolacyltransferase (Dgat1), perilipin (Plin1), mitoguardin 2 (Miga2), and poly(ADP-ribose) polymerase1(Parp1). Results are presented as mean ± SD of one representative out of three independent experiments. Normalization was done by taking geometric mean of three housekeeping genes, Importin, Tubulin and AcTH.

