## Supplementary material for "High-energy demand and nutrient exhaustion in MTCH2 knockout cells": Table1

### Table1_Heatmap_Figure.2SD

| S.no | Metabolites | WT.MKO.fc | WT.MKO.pval | WT.MKO.fdr | WT.MKO-R.fc | WT.MKO-R.pval | WT.MKO-R.fdr |
|---|---|---|---|---|---|---|---|
| 1 | 1-Deoxy-D-xylulose-5-P (DXP) | -1.592 | 0.005 | 0.016 | -1.327 | 0.014 | 0.035 |
| 2 | 2'-Deoxycytidine | -6.156 | 0.006 | 0.017 | 1.135 | 0.698 | 0.796 |
| 3 | 2'-Deoxyguanosine | -3.024 | 0.002 | 0.009 | 1.266 | 0.146 | 0.235 |
| 4 | 2-Hydroxy-3-methylbutyric | -1.945 | 0.000 | 0.003 | -0.379 | 0.424 | 0.549 |
| 5 | 2-Hydroxybutyrate | -2.089 | 0.003 | 0.010 | 0.073 | 0.982 | 0.999 |
| 6 | 2-Hydroxyoctanoic acid | -1.979 | 0.000 | 0.002 | -1.394 | 0.002 | 0.007 |
| 7 | 2-Quinolinecarboxylic acid | -3.073 | 0.000 | 0.000 | -3.134 | 0.000 | 0.000 |
| 8 | 3-Methyladenine | -2.098 | 0.006 | 0.017 | -0.569 | 0.481 | 0.610 |
| 9 | 3-Sulphino-L-alanine | -1.812 | 0.003 | 0.010 | -1.963 | 0.002 | 0.006 |
| 10 | 4-Hydroxyphenyllactic acid | -2.573 | 0.000 | 0.003 | 2.337 | 0.001 | 0.003 |
| 11 | 5-Keto-D-gluconic acid | -2.465 | 0.001 | 0.004 | -2.893 | 0.000 | 0.001 |
| 12 | 6-Phosphogluconic acid | -2.633 | 0.000 | 0.003 | -2.854 | 0.000 | 0.001 |
| 13 | Adenine | -2.580 | 0.000 | 0.003 | -5.091 | 0.000 | 0.000 |
| 14 | Adenosine | -3.177 | 0.003 | 0.010 | -4.407 | 0.000 | 0.001 |
| 15 | 3'-AMP | -1.415 | 0.006 | 0.017 | -0.103 | 0.940 | 0.983 |
| 16 | Alpha-Hydroxyglutaric acid | -2.080 | 0.001 | 0.005 | 1.424 | 0.010 | 0.027 |
| 17 | Alpha-Ketoglutaric acid | -2.559 | 0.006 | 0.016 | -0.210 | 0.924 | 0.975 |
| 18 | Benzoate | -2.208 | 0.001 | 0.004 | -2.945 | 0.000 | 0.001 |
| 19 | Beta-Hydroxyisovaleric acid | -1.567 | 0.005 | 0.015 | 0.667 | 0.189 | 0.287 |
| 20 | Citric acid | -3.076 | 0.001 | 0.005 | -3.129 | 0.001 | 0.004 |
| 21 | Citrulline | -2.237 | 0.003 | 0.010 | -2.117 | 0.004 | 0.012 |
| 22 | D-galactonate | -2.272 | 0.000 | 0.003 | -2.602 | 0.000 | 0.001 |
| 23 | D-Glucosamine 6-phosphate | -3.892 | 0.001 | 0.005 | -1.878 | 0.057 | 0.115 |
| 24 | Dodecyl sulfate | -3.184 | 0.000 | 0.001 | -2.295 | 0.001 | 0.004 |
| 25 | D-Ribose 5-phosphate | -2.191 | 0.000 | 0.000 | -0.493 | 0.147 | 0.235 |
| 26 | Fumaric acid | -2.547 | 0.000 | 0.000 | 0.140 | 0.851 | 0.917 |
| 27 | GABA | -1.942 | 0.000 | 0.002 | -0.301 | 0.519 | 0.637 |
| 28 | Glucose 6-phosphate | -1.967 | 0.000 | 0.000 | -2.270 | 0.000 | 0.000 |
| 29 | Glutamic acid | -1.901 | 0.000 | 0.003 | -0.217 | 0.728 | 0.821 |
| 30 | Glyceraldehyde 3-phosphate | -1.476 | 0.000 | 0.003 | -0.751 | 0.027 | 0.060 |

| | | | | | | | |
|---|---|---|---|---|---|---|---|
| 31 | Glycerol | -1.909 | 0.001 | 0.007 | -0.935 | 0.066 | 0.128 |
| 32 | Guanine | -2.475 | 0.001 | 0.005 | 4.672 | 0.000 | 0.000 |
| 33 | Guanosine | -1.722 | 0.004 | 0.012 | 2.779 | 0.000 | 0.001 |
| 34 | Hexanoic acid | -2.312 | 0.000 | 0.001 | -2.602 | 0.000 | 0.000 |
| 35 | Leu-Ala | -2.109 | 0.004 | 0.013 | -2.205 | 0.003 | 0.010 |
| 36 | Levulinic acid | -2.144 | 0.000 | 0.000 | -2.198 | 0.000 | 0.000 |
| 37 | Lumichrome | -2.302 | 0.000 | 0.002 | -1.104 | 0.020 | 0.045 |
| 38 | Malic acid | -2.471 | 0.000 | 0.000 | 0.032 | 0.990 | 0.999 |
| 39 | Mannose 6-phosphate | -2.211 | 0.000 | 0.000 | -1.045 | 0.003 | 0.010 |
| 40 | Mesaconic acid | -1.753 | 0.002 | 0.007 | 0.726 | 0.126 | 0.205 |
| 41 | Methionine sulfoxide | -1.966 | 0.002 | 0.009 | -0.265 | 0.763 | 0.846 |
| 42 | Methyl vanillate | -2.069 | 0.000 | 0.003 | -1.735 | 0.002 | 0.006 |
| 43 | Myo-Inositol | -1.727 | 0.004 | 0.013 | 2.441 | 0.000 | 0.002 |
| 44 | N-Acetyl-L-methionine | -2.334 | 0.002 | 0.009 | -1.109 | 0.092 | 0.166 |
| 45 | N-Formyl-L-Methionine | -1.755 | 0.003 | 0.012 | -0.917 | 0.087 | 0.159 |
| 46 | O-Phosphoethanolamine | -2.038 | 0.000 | 0.000 | -0.316 | 0.333 | 0.452 |
| 47 | Oxalate | -2.390 | 0.000 | 0.000 | -1.855 | 0.000 | 0.000 |
| 48 | Phe-Ala | -2.383 | 0.004 | 0.013 | -2.574 | 0.003 | 0.009 |
| 49 | Phe-Gly | -3.067 | 0.005 | 0.016 | -3.098 | 0.005 | 0.014 |
| 50 | Phenol red | -1.450 | 0.001 | 0.004 | -1.753 | 0.000 | 0.001 |
| 51 | Phenylacetic acid | -2.049 | 0.001 | 0.004 | -1.318 | 0.011 | 0.027 |
| 52 | Phenylpyruvic acid | -2.784 | 0.002 | 0.007 | -2.209 | 0.007 | 0.019 |
| 53 | Phosphoenolpyruvate | -1.931 | 0.003 | 0.011 | -4.619 | 0.000 | 0.000 |
| 54 | Phosphoric acid | -1.886 | 0.000 | 0.001 | -1.414 | 0.001 | 0.004 |
| 55 | Phosphorylcholine | -1.781 | 0.002 | 0.007 | 0.154 | 0.879 | 0.942 |
| 56 | Phthalic acid | -2.199 | 0.000 | 0.000 | -1.994 | 0.000 | 0.000 |
| 57 | Pro-Gly | -3.153 | 0.000 | 0.002 | -3.658 | 0.000 | 0.000 |
| 58 | Pyrrole-2-carboxylic acid | -2.246 | 0.002 | 0.009 | -1.463 | 0.027 | 0.059 |
| 59 | Riboflavin | -1.802 | 0.002 | 0.007 | -2.578 | 0.000 | 0.001 |
| 60 | Saccharate | -1.476 | 0.000 | 0.002 | -2.779 | 0.000 | 0.000 |
| 61 | S-Adenosylhomocysteine | -1.641 | 0.000 | 0.003 | -0.793 | 0.035 | 0.072 |
| 62 | Sedoheptulose 7-phosphate | -2.214 | 0.000 | 0.002 | -1.375 | 0.005 | 0.014 |
| 63 | S-Hexylglutathione | -3.024 | 0.001 | 0.005 | -2.982 | 0.001 | 0.005 |
| 64 | Sucrose | -5.513 | 0.001 | 0.005 | -18.263 | 0.000 | 0.000 |

| | | | | | | | |
|---|---|---|---|---|---|---|---|
| 65 | Tartrate | -2.462 | 0.001 | 0.006 | -1.984 | 0.005 | 0.014 |
| 66 | Thiolactic acid | -2.359 | 0.000 | 0.000 | -2.100 | 0.000 | 0.000 |
| 67 | Threonine | -2.165 | 0.006 | 0.016 | -0.888 | 0.225 | 0.329 |
| 68 | Thymidine | -2.826 | 0.005 | 0.016 | -0.354 | 0.834 | 0.906 |
| 69 | Thymine | -2.679 | 0.006 | 0.018 | -0.234 | 0.918 | 0.975 |
| 70 | Vanillin | -2.796 | 0.001 | 0.004 | -3.834 | 0.000 | 0.001 |
