## Supplementary material for "High-energy demand and nutrient exhaustion in MTCH2 knockout cells": Table2

### Table2_Heatmap_Figure.3A

| Groups | p.MKO_WT | p.WT_MKO-R | p.KO_MKO-R | fdr | fdr.MKO_WT | p.MKO_WT | p.WT_MKO-R | p.KO_MKO-R | fdr | fdr.MKO_WT |
|---|---|---|---|---|---|---|---|---|---|---|
| FC | 1.837E-06 | 4.686E-01 | 4.507E-06 | 4.992E-06 | 4.042E-05 | -4.682E-02 | -9.367E-02 | -1.405E-01 | -1.874E-01 | -2.342E-01 |
| TC | 1.162E-05 | 1.673E-02 | 3.838E-04 | 2.994E-05 | 6.596E-05 | -1.533E-03 | -3.193E-03 | -4.852E-03 | -6.511E-03 | -8.171E-03 |
| FA | 3.640E-03 | 4.509E-03 | 2.460E-05 | 5.005E-05 | 5.719E-03 | 2.699E-03 | 2.669E-03 | 2.639E-03 | 2.609E-03 | 2.579E-03 |
| FFA | 1.405E-04 | 1.141E-03 | 2.337E-01 | 1.929E-04 | 3.434E-04 | 4.695E-02 | 4.689E-02 | 4.684E-02 | 4.678E-02 | 4.673E-02 |
| CE | 1.092E-05 | 2.409E-09 | 8.126E-10 | 3.257E-09 | 6.596E-05 | 4.840E-05 | 5.941E-05 | 7.041E-05 | 8.142E-05 | 9.243E-05 |
| DAG | 2.472E-03 | 2.476E-04 | 3.054E-06 | 1.137E-05 | 4.356E-03 | 2.478E-03 | 2.831E-03 | 3.184E-03 | 3.537E-03 | 3.890E-03 |
| TAG | 1.499E-05 | 4.923E-03 | 6.352E-07 | 4.339E-06 | 6.596E-05 | -4.433E-04 | -9.250E-04 | -1.407E-03 | -1.888E-03 | -2.370E-03 |
| CL | 1.297E-05 | 2.110E-03 | 2.832E-03 | 2.994E-05 | 6.596E-05 | 4.180E-04 | 2.206E-04 | 2.322E-05 | -1.742E-04 | -3.715E-04 |
| LCL | 1.053E-03 | 1.159E-04 | 1.202E-06 | 5.998E-06 | 2.107E-03 | 1.256E-03 | 1.455E-03 | 1.655E-03 | 1.855E-03 | 2.055E-03 |
| LPA | 8.175E-01 | 9.565E-01 | 9.432E-01 | 8.323E-01 | 8.175E-01 | 8.362E-01 | 8.238E-01 | 8.113E-01 | 7.989E-01 | 7.865E-01 |
| LPC | 1.322E-01 | 1.381E-04 | 1.687E-05 | 2.994E-05 | 1.530E-01 | 6.956E-02 | 7.373E-02 | 7.789E-02 | 8.205E-02 | 8.622E-02 |
| LPE | 5.875E-02 | 3.023E-02 | 6.759E-04 | 1.045E-03 | 7.603E-02 | 3.496E-02 | 3.550E-02 | 3.604E-02 | 3.657E-02 | 3.711E-02 |
| LPG | 3.421E-04 | 2.984E-03 | 2.596E-01 | 4.365E-04 | 7.527E-04 | 5.230E-02 | 5.213E-02 | 5.196E-02 | 5.178E-02 | 5.161E-02 |
| LPI | 7.906E-03 | 7.964E-04 | 1.265E-05 | 2.994E-05 | 1.160E-02 | 6.052E-03 | 6.713E-03 | 7.374E-03 | 8.035E-03 | 8.696E-03 |
| LPS | 3.219E-01 | 1.411E-01 | 1.328E-02 | 1.718E-02 | 3.540E-01 | 1.516E-01 | 1.457E-01 | 1.397E-01 | 1.338E-01 | 1.278E-01 |
| PA | 4.039E-01 | 2.433E-04 | 1.264E-03 | 2.815E-04 | 4.232E-01 | 1.773E-01 | 1.812E-01 | 1.850E-01 | 1.889E-01 | 1.927E-01 |
| PC | 1.172E-01 | 1.338E-05 | 2.311E-06 | 6.099E-06 | 1.433E-01 | 6.772E-02 | 7.293E-02 | 7.814E-02 | 8.335E-02 | 8.856E-02 |
| PE | 2.824E-05 | 2.243E-04 | 2.363E-07 | 3.466E-06 | 8.876E-05 | 3.906E-05 | 2.907E-05 | 1.909E-05 | 9.108E-06 | -8.746E-07 |
| PG | 1.881E-05 | 1.926E-04 | 1.005E-01 | 2.994E-05 | 6.896E-05 | 2.013E-02 | 2.013E-02 | 2.012E-02 | 2.011E-02 | 2.011E-02 |
| PI | 3.730E-02 | 2.814E-04 | 1.442E-05 | 2.994E-05 | 5.129E-02 | 2.610E-02 | 2.888E-02 | 3.165E-02 | 3.442E-02 | 3.719E-02 |
| PS | 1.049E-04 | 2.821E-04 | 5.706E-07 | 4.339E-06 | 2.884E-04 | 1.628E-04 | 1.718E-04 | 1.807E-04 | 1.896E-04 | 1.985E-04 |
| SM | 2.574E-03 | 9.355E-01 | 1.605E-03 | 1.088E-03 | 4.356E-03 | -9.023E-02 | -1.833E-01 | -2.764E-01 | -3.695E-01 | -4.626E-01 |
