## Supplementary material for "High-energy demand and nutrient exhaustion in MTCH2 knockout cells": Table3

**Table.3_List of Primers_Fig 4B and Fig 4SB**

| S.no | Gene | Primers | Sequences |
|---|---|---|---|
| 1 | Fabp4 | F | AAGTGGGAGTGGGCTTTGC |
| | | R | CCGGATGGTGACCAAATCC |
| 2 | Parp1 | F | TGGTTTCAAGTCCCTTGTCC |
| | | R | TGCTGTCTATGGAGCTGTGG |
| 3 | Fasn | F | GGAGGTGGTGATAGCCGGTAT |
| | | R | TGGGTAATCCATAGAGCCCAG |
| 4 | Pdha1 | F | GAAATGTGACCTTCATCGGCT |
| | | R | TGATCCGCCTTTAGCTCCATC |
| 5 | Scd1 | F | TTCTTGCGATACACTCTGGTGC |
| | | R | CGGGATTGAATGTTCTTGTCGT |
| 6 | Fam73b(Miga2) | F | GGAGGACTGAGGGTATGTCCA |
| | | R | CAAGGGCTGTGGCAAAAAGA |
| 7 | cfd/Adipsin | F | CATGCTCGGCCCTACATGG |
| | | R | CACAGAGTCGTCATCCGTCAC |
| 8 | AGPAT | F | TAAGATGGCCTTCTACAACGGC |
| | | R | CCATACAGGTATTTGACGTGGAG |
| 9 | DGAT1 | F | TCCGTCCAGGGTGGTAGTG |
| | | R | TGAACAAAGAATCTTGCAGACGA |
| 10 | AdipoQ | F | GACAAGGCCGTTCTCTTCAC |
| | | R | CAGACTTGGTCTCCCACCTC |
| 11 | Cebpa | F | GAACAGCAACGAGTACCGGGTA |
| | | R | GCCATGGCCTTGACCAAGGAG |
| 12 | Cebpb | F | CAAGCTGAGCGACGAGTACA |
| | | R | CAGCTGCTCCACCTTCTTCT |
| 13 | Cebpd | F | TGCCCACCCTAGAGCTGTG |
| | | R | CGCTTTGTGGTTGCTGTTGA |
| 14 | Pparg2 | F | TGCTGTTATGGGTGAAACTCT |
| | | R | CGCTTGATGTCAAAGGAATGC |
| 15 | PLIN1 | F | GGGACCTGTGAGTGCTTCC |
| | | R | GTATTGAAGAGCCGGGATCTTTT |
| 16 | Importin8 | F | CAGCAGGATTGCTTCGAGTA |
| | | R | AGCATAGCACTCGGCATCTT |
| 17 | mTubulin | F | CCAGGGCTTCTTGGTTTTCC |
| | | R | CGCTCAATGTCGAGGTTTCT |
| 18 | AcTH | F | CCTGATCCACATCTGCTGGAA |
| | | R | ATTGCCGACAGGATGCAGAA |
